# M1 macrophage-mediated lymphangiogenesis aggravates liver fibrosis via MDK/YAP signaling pathway

**DOI:** 10.64898/2026.05.25.727562

**Authors:** Dan Wang, Dan Long, Ying Zhao, Dian Li, Fenglin Xiong, Ziwei Huang, Liping Yang, Qing Zheng, Yonghua Chen, Yanni Zhou, Li Feng

## Abstract

**Background:** Lymphangiogenesis plays a critical role in various liver diseases, yet its function in liver fibrosis remains controversial. This study aimed to explore the role of lymphangiogenesis in liver fibrogenesis and its underlying regulatory mechanisms.

**Methods:** Liver fibrotic mice were established by carbon tetrachloride (CCl_4_) or Thioacetamide (TAA)-induced injection or bile duct ligation. Lymphatic vessels were marked by podoplain (Pdpn) staining in mice and D2-40 staining in clinical samples. Lymphatic vessels area and density were measured to indicate lymphangiogenesis. Multiplexing immunohistochemistry was used to detect co-localization of proteins.

**Results:** In the present study, we first verified increased lymphangiogenesis in human and murine fibrotic livers. Afterwards, we identified VEGFC rather than VEGFD as the primary driver of lymphangiogenesis in liver fibrosis. Furthermore, we demonstrated that M1 macrophages serve as the major source of VEGFC. Founctional studies revealed that VEGFC-mediated lymphangiogenesis exacerbates hepatic fibrosis, while its inhibition alleviated fibrosis. Bioinformatic analysis uncovered Midkine (MDK) as a key downstream of lymphangiogenesis. Both *in vivo* and *in vitro* studies confirmed that exogenous MDK promotes liver fibrosis via activating hepatic stellate cells (HSCs), whereas MDK inhibition counteracts the profibrotic effects of VEGFC-induced lymphangiogenesis. Importantly, we discovered that MDK activates HSCs through the Hippo/YAP signaling pathway.

**Conclusions:** M1 macrophage-mediated lymphangiogenesis aggravates liver fibrosis via MDK secretion, which activates HSCs. These findings provide novel insights into coordinated crosstalk between macrophages, lymphatic endothelial cells and HSCs in liver fibrosis and suggest lymphangiogenesis and MDK as potential therapeutic targets for fibrotic liver diseases.

## 1 Background

Liver fibrosis is the result of persistent liver injury caused by various etiological factors, leading to excessive deposition of extracellular matrix (ECM). It is an essential stage in the progression of chronic liver disease to cirrhosis and hepatocellular carcinoma (HCC). The lymphatic system plays a crucial role in numerous vital physiological processes within the body, including tissue fluid homeostasis and immune cell transport. The liver, as the largest lymph-producing organ, generates 25-50% of the body’s total lymph volume ^[1]^. The accumulation of lymphatic fluid leading to ascites due to lymphatic system dysfunction is a typical clinical manifestation in patients with liver fibrosis and cirrhosis. Lymphangiogenesis, characterized by an increase in lymphatic vessel density (LVD) and/or luminal area, has been reported to play a significant role in the pathogenesis of various liver diseases. For instance, platelet-derived growth factor-D (PDGF-D) has been shown to promote lymph node metastasis in cholangiocarcinoma by enhancing lymphangiogenesis ^[2]^. In both hepatocellular carcinoma and intrahepatic cholangiocarcinoma, lymphangiogenesis is associated with poor prognosis ^[3]^. Conversely, during acute cellular rejection (ACR) in liver transplantation, lymphangiogenesis coincides with the attenuation of ACR and inflammation, as well as prolonged graft survival ^[3, 4]^.

Beyond its established roles in pathological processes such as tumor metastasis and liver transplantation, lymphangiogenesis has also been implicated in the progression of liver fibrosis and cirrhosis. Tanaka and Jeong et al. independently reported increased lymphangiogenesis in various experimental models, including: partial portal vein ligation and bile duct ligation (BDL) in rat fibrosis models, BDL and carbon tetrachloride (CCl_4_)-induced mouse fibrosis models, as well as in human cirrhosis patients ^[5, 6]^. Utilizing single-cell RNA sequencing, Su et al. discovered a remarkable 20-fold increase in the proportion of lymphatic endothelial cells (LECs) among total endothelial cells in CCl_4_-induced mouse liver fibrosis models ^[7]^. These collective findings strongly suggest that lymphangiogenesis is augmented during liver fibrosis progression.

However, the role of lymphangiogenesis in liver fibrosis and cirrhosis remains poorly understood, with limited studies and conflicting conclusions in existing researches. It’s reported that in BDL-induced liver fibrosis models in rats, sympathetic nerves promote lymphangiogenesis, while fibrosis is exacerbated following sympathetic denervation. The authors therefore propose that lymphangiogenesis may exert anti-fibrotic effects. However, they acknowledge the possibility that sympathetic denervation might aggravate fibrosis by enhancing ductular reaction rather than through suppressing lymphangiogenesis ^[5]^. Another study found that acute-on-chronic liver failure exhibits more severe hepatic injury and inflammation than cirrhosis, accompanied by lymphatic vessel reduction due to LEC apoptosis ^[7]^. Additional research demonstrates that in long-term high-fat/high-cholesterol diet-induced fibrotic models, while lymphangiogenesis increases, the expression of lymphatic markers PDPN, LYVE1 and PROX1 is downregulated with concomitant impairment of lymphatic drainage function. VEGFC-induced lymphangiogenesis restored drainage capacity and partially alleviated inflammation, though without significant improvement in ALT/AST levels ^[8]^. These findings collectively suggest that lymphangiogenesis may help mitigate liver injury and fibrosis progression, while other studies indicate opposing effects. Huang et al. identified through transcriptome sequencing that LECs in CCl_4_- and BDL-induced fibrotic livers highly express various collagens including col1a1 and col1a2 ^[9]^. However, it remains unclear whether these collagens exacerbate ECM deposition or modify lymphatic permeability through basement membrane remodeling. Another investigation revealed that compared to healthy human livers, end-stage livers from non-alcoholic steatohepatitis (NASH) and HCV progression showed increased lymphatic density, with LECs overexpressing CCL21 and surrounded by abundant macrophages and T cells. The authors propose this lymphangiogenesis may represent a host response to inflammatory cell infiltration. Single-cell sequencing further identified activated IL13 signaling in LECs causing PROX1 downregulation, potentially increasing lymphatic permeability and contributing to ascites formation ^[10]^. Hence, more conclusive evidence is required to clarify the role of lymphangiogenesis in liver fibrogenesis.

In this study, we demonstrated that VEGFC—rather than VEGFD—is the pivotal cytokine driving lymphangiogenesis in liver fibrosis and revealed its primary source as M1-polarized macrophages. We then regulated lymphangiogenesis through VEGFC administration and inhibition to evaluate its impact on liver fibrosis. To uncover the mechanism by which lymphangiogenesis exacerbates fibrosis, we performed bioinformatic analysis of single-cell RNA sequencing (scRNA-seq) data, identifying midkine (MDK)—a factor secreted by LECs—as a key mediator of crosstalk between lymphangiogenesis and hepatic stellate cells (HSCs).

## 2 Methods

### 2.1 Clinical samples

Clinical tissue samples were obtained from Biobank of West China Hospital. Written informed consent was obtained from participants before tissue collection, and all tissue samples were freshly frozen in liquid nitrogen and stored at −80 °C. The Ethics Committee of Sichuan University approved the use of the clinical samples and the study.

### 2.2 Animal experiments

The Committee on Animal Research of Sichuan University approved all animal experiments. All animal studies were conducted in accordance with the “Guidelines for the Care and Use of Laboratory Animals of the National Institutes of Health”. Eight-week-old C57BL/6J male mice were purchased from Beijing Vital River Laboratory Animal Technology (Beijing, China). All mice were housed in separate cages with 12 h light-dark cycles at a temperature of 25 °C and a humidity of 60∼65 %. To induce liver fibrosis, the mice were injected intraperitoneally twice a week with 5% CCl_4_ (Merck, Germany; 5 μL/g body weight in olive oil). Mice were randomly assigned to corresponding groups after two weeks of CCl_4_ injection. Afterwards, mice were injected with VEGFC and/or MAZ-51 or iMDK. VEGFC (Yeasen biotechnology, Shanghai, China; 0.5 μg/g body weight in normal saline) were injected through tail vein every other day. MAZ-51 (MedChemExpress, Shanghai, China; 8 mg/kg) or iMDK (MedChemExpress; 8 mg/kg) were injected intraperitoneally twice a week. For BDL model, bile ducts were double ligated and specimens were collected at the 3rd week. Further treatment was applied from the 2^nd^ week and last 2 weeks. For the liver fibrosis remission model, mice were intraperitoneally injected with 10% CCl_4_ for six consecutive weeks, followed by a two-week drug withdrawal period. For VEGFC group, CCl_4_ administration was discontinued and simultaneously replaced with VEGFC treatment. For Thioacetamide-induced (TAA; MedChemExpress; 8 mg/kg) liver fibrosis, mice were injected intraperitoneally twice a week for 8 weeks.

The mice were euthanized at the end of the treatment period, and samples were collected. Dead mice during treatment will be excluded from the analysis. Serum levels of alanine transaminase (ALT), aspartate aminotransferase (AST) and Hydroxyproline content were measured according to the manufacture’s instruction using commercial kits, purchased from Nanjing Jiancheng Bioengineering Institute (Nanjing, China).

### 2.3 Cell cultures and tubule formation experiment

Human primary lymphatic endothelial cell line was purchased from Kewen Biotechnology (Wuhan, China) and cultured in specific culture media purchased from the same company. LX-2, a gift from Dr. Xiaofeng Lu, was cultured in Dulbecco’s modified Eagle’s medium (Gibco, NY, USA), supplied with 10% fetal bovine serum (Merck, Germany). When present, MDK, purchased from MedChemExpress and Verteporfin, purchased from Macklin (Shanghai, China), were added to culture medium at the indicated concentration. Thp-1, a gift from Dr. Jinnan Li, was cultured in RPMI media 1640 (Gibco, NY, USA) supplied with 10% fetal bovine serum (Merck). For macrophage induction, Phorbol myristate acetate (PMA), purchased from MedChemExpress, was added into culture medium at a final concentration of 100 ng/mL. To induce M1 polarization, IFN-γ (Yeasen, Shanghai, China) and LPS (MedChemExpress) were added at a final concentration of 20 ng/mL and 100 ng/mL, respectively. To induce M2 polarization, IL4 and IL13, purchased from NovoProtein Scientific Inc. (Suzhou, China), were added at a final concentration of 20 ng/mL.

For tubule formation experiment, 96 well-plate was covered by Matrigel (Corning, NY, USA), and then 1000 LECs were seeded, with/without VEGFC or VEGFD (NovoProtein, Suzhou, China) added at a final concentration of 10ng/mL. Tubule formation was detected after 24 h treatment.

### 2.4 Determination of lymphatic vessel density (LVD) and lymphatic vessel area (LVA)

Lymphangiogenesis in liver tissues was estimated by calculating LVD and LVA according to established methods as described previously ^[11]^. Briefly, lymphatic vessels were stained with anti-D2-40 in human tissues and with anti-Pdpn in murine tissues. Individual lymphatic vessels around portal veins were counted and number of lymphatic vessels around each portal vein was calculated as LVD. At least four areas were randomly selected in each sample and LVA was measured.

### 2.5 Immunofluorescence

Paraffin sections were routinely dewaxed and hydrated, antigens were recovered by a microwave □ based antigen retrieval method. Primary antibodies against Pdpn (AF3244, R&D), VEGFR3 (AF743, R&D), Lyve1 (ab218535, Abcam), Ki67 (A23722, Abclonal), VEGFC (YT5297, Immunoway) and F4/80 (70076, CST) were incubated at 4 °C overnight. Secondary antibodies conjugated with Alexa Fluor® 488 (ab150077, Abcam), 555 (A-21428, Invitrogen) or 647 (ab150135, Abcam) were added. The results were visualized using a VS200 Whole Slide Fluorescence Scanning System or NIKON confocal microscope. Images were analyzed by corresponding softwares and Image J.

### 2.6 Immunohistochemistry and multiplexing immunohistochemistry (mIHC)

To analyze the expression level of α-sma and VEGFC and mark lymphatic vessels, immunohistochemistry was conducted on 5 µm FFPE sections of human and murine liver tissues. A microwave □based antigen retrieval method was employed to expose and unmask the target antigens in the tissue sections. The samples were then incubated with primary antibodies against α-sma (ET1607-53, HuaBio), VEGFC (YT5297, Immunoway) or D2-40 (ZM-0465, ZSGB-Bio). Subsequently, a secondary antibody conjugated to horseradish peroxidase (HRP, Jackson) was added, producing a visible signal with the substrate 3,3′□diaminobenzidine (DAB, CST). The results were visualized using a VS200 Whole Slide Brightfield Scanning System and analyzed by corresponding software.

For multiplexing immunohistochemistry, Mouse/Rabbit Triple-Target Four-Color Fluorescence Detection Kit (RS0035, Immunoway) was used and experiments were conducted according to the protocol provided by the manufacture. Antibodies against F4/80 (70076, CST), CD206 (AF2353-SP, R&D), CD68 (ab303565, Abcam), CD86 (19589S, CST), CD3 (78588S, CST) and VEGFC (YT5297, Immunoway) were used.

### 2.7 Immunoblotting

Total proteins were extracted and concentration was quantified by a BCA protein assay kit (Thermo Fisher, MA, USA). 10-20□µg protein was loaded, separated on 10% or 12% SDS-PAGE, and transferred to PVDF membranes. Then membranes were blotted at 4□°C overnight with primary antibodies against α-sma (ET1607-53, HuaBio), collagen I (HA722517, HuaBio), YAP (14074, CST), pYAP (13008, CST), MOB1 (13730, CST), MST1 (3682, CST), MDK (11009-1-AP, Proteintech), VEGFC (YT5297, Immunoway), PROX1 (ab199359, Abcam), iNOS (ER1706-89, Huabio), CD206 (YT5640, Immunoway), β-actin (R380624, Zenbio) and β-Tubulin (TA-10, ZSGB-Bio). Subsequently, a secondary antibody conjugated to HRP (Jackson) was added. The results were visualized using Fusion X (France). For analysis, expression of proteins were normalized to β-actin or β-Tubulin.

### 2.8 Enzyme-linked immunosorbent assays

The level of IL-6, IL-1β and TGF-β1 in liver tissues and VEGFC and MDK in serum was measured using enzyme-linked immunosorbent assay kit (Kamaishu biotechology, Shanghai, China). Liver tissues were homogenized in cold radioimmunoprecipitation assay buffer (Beyotime, Jiangsu, China) supplemented with a 1% proteinase inhibitor cocktail (Yeasen, China). The protein concentration of the samples was determined, followed by target proteins concentration detection using the kits.

### 2.9 Real-time quantitative polymerase chain reaction analysis

Total RNA was extracted from liver tissues or cells using Total RNA extraction kit (Promega, WI, USA) according to the manufacture’s protocol. One microgram of RNA was reverse-transcribed using the First Strand cDNA Synthesis Kit (Yeasen, China). qPCR was performed using SYBR Master Mix (Yeasen, China) and analyzed by QuantStudio 6 Flex Real-Time PCR System (Thermo Fisher, USA). mRNA expression levels were calculated using the 2−^ΔΔ^Ct approach and normalized to RPS18. The primer sequences were listed in Table S1.

### 2.10 Statistics

Quantitative data shown are representative of at least three separate experiments and expressed as means ± the standard deviation of the mean. For comparisons between the two groups, unpaired Student’s t-test was used. For comparisons among three or more groups, one-way or two-way analysis of variance was used. Statistical analysis was performed using GraphPad software (version 9.0, GraphPad Software, La Jolla, CA, USA). P < 0.05 was considered statistically significant.

## 3 Results and discussion

### 3.1 Lymphangiogenesis is associated with liver fibrogenesis

To elucidate changes in lymphatic vessels during liver fibrogenesis, we first conducted scRNA-seq analysis with a public cirrhosis dataset GSE136103 and found increasing LECs in cirrhotic patients compared to healthy controls (Fig. 1a-c). We next examined lymphangiogenesis in cirrhotic human liver tissues by D2-40 immunohistochemistry. Quantitative analysis revealed significantly increased LVD and LVA surrounding portal veins compared to controls (Fig. 1d-f). Moreover, lymphangiogenesis gradually increased along with fibrosis progression (Fig. 1g-i). To validate these findings, we analyzed lymphatic vessel formation (PDPN staining) in multiple murine fibrosis models, including CCl_4_-, BDL-, TAA- and DDC-induced fibrosis. Consistent with human data, all fibrotic models exhibited marked lymphangiogenesis around portal areas (Fig. 1j-r and Supplementary Fig. 1).

**Fig. 1.**
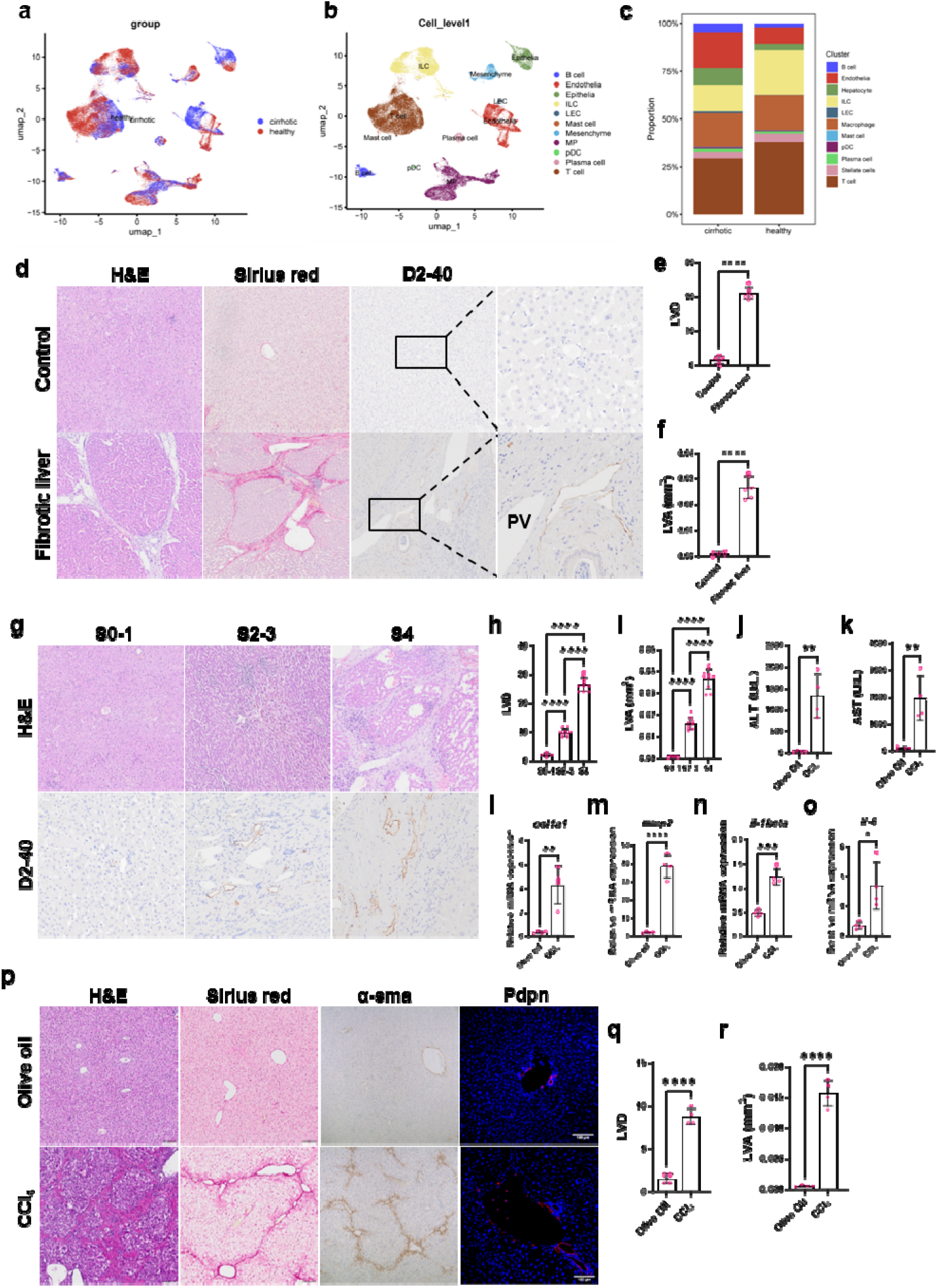
Increased lymphangiogenesis in fibrotic livers. **a-b** UMAP plot of non-parenchymal cells in single-cell RNA sequencing of liver cirrhosis samples. **c** Analysis of proportion changes in liver LECs in cirrhotic tissues. **d** Control and cirrhotic liver samples stained with H&E, sirius red and D2-40. **e-f** Calculation of LVD (**e**) and LVA (**f**) by statistical analysis of D2-40 staining (n=5-6/group). **g** Liver tissue samples at different fibrosis stages stained with H&E and D2-40. **h-i** Calculation of LVD (**h**) and LVA (**i**) by statistical analysis of D2-40 staining (n=5-10/group). **j-k** Serum ALT (**j**) and AST (**k**) levels were measured. **l-o**. mRNA levels of *col1a1* (**l**), *mmp2* (**m**), *il-1beta* (**n**) and *il-6* (**o**) were quantified by RT-qPCR. **p** Pathological section examination and IHC staining of lymphatic vessel marker-pdpn. **q-r** Calculation of LVD (**q**) and LVA (**r**) by statistical analysis of pdpn staining. N=4/group. *, P<0.05; **, P<0.01; ***, P<0.001; ****, P<0.0001.

To determine whether lymphangiogenesis reverses during fibrosis regression, we analyzed livers after cessation of CCl_4_ treatment. Histological assessment demonstrated that lymphatic vessel formation decreased in parallel with fibrosis resolution (Supplementary Fig. 2). These results establish a dynamic correlation between lymphangiogenesis and liver fibrogenesis.

### 3.2 VEGFC, but not VEGFD, drives fibrotic lymphangiogenesis

VEGFC and VEGFD are classic cytokines to induce lymphangiogenesis. Surprisingly, we found that only VEGFC administration promoted lymphangiogenesis *in vivo*, while VEGFD failed (Fig. 2a-c). This specificity was confirmed *in vitro*, where VEGFC (but not VEGFD) promoted tube formation in primary LECs (Fig. 2d). Besides, transcriptome analysis revealed that VEGFC treatment significantly enriched pathways associated with tube morphogenesis in LECs (Supplementary Fig. 3). In CCl_4_- and BDL-induced murine fibrosis models, increased VEGFC expression in mRNA and protein levels was detected (Fig. 2e-g). Furthermore, clinical relevance was supported by elevated serum and tissue VEGFC levels in cirrhotic patients (Fig. 2h-i).

**Fig. 2.**
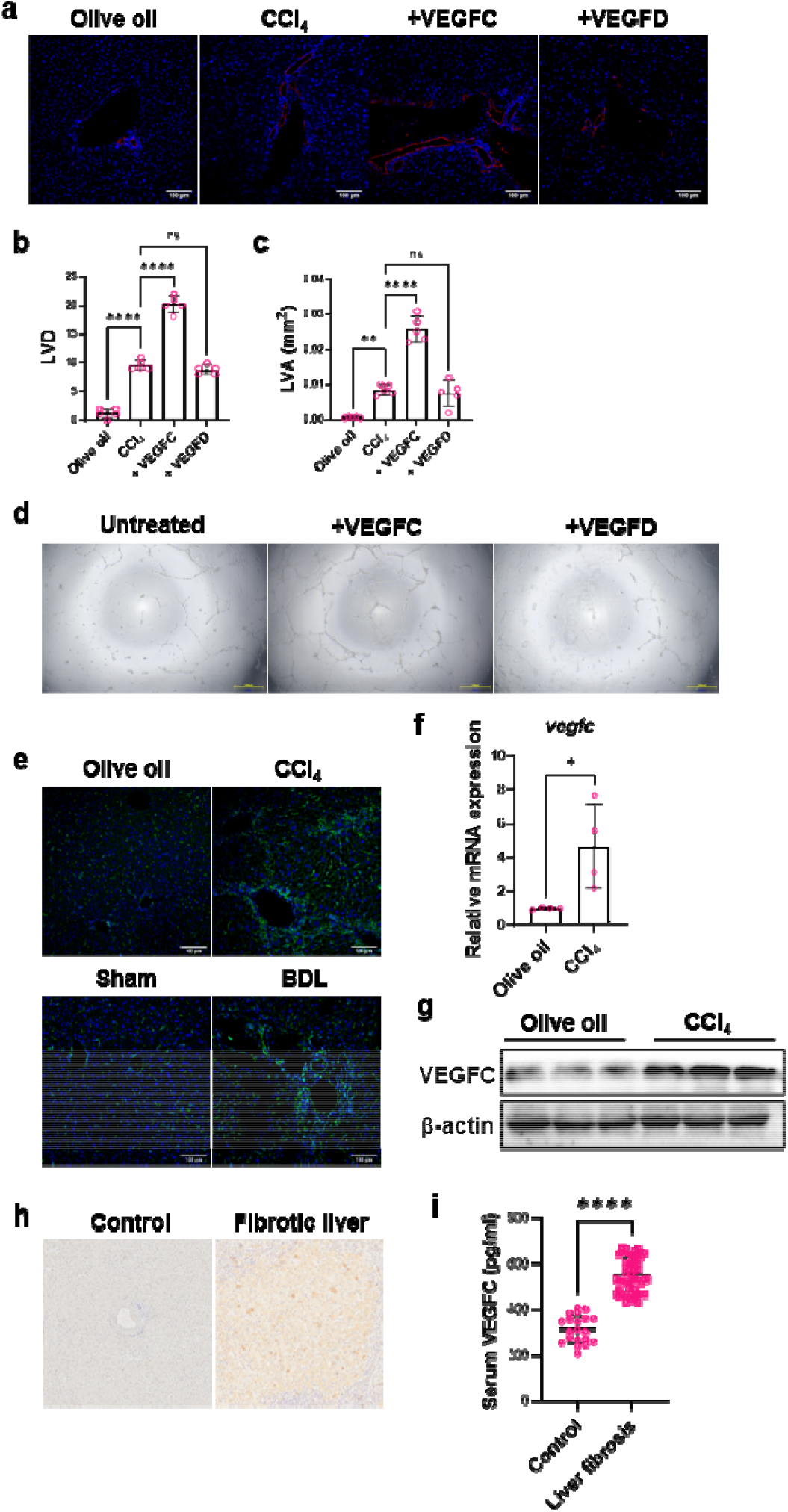
VEGFC, instead of VEGFD induces lymphangiogenesis. **a** Lymphatic vessels were marked by Pdpn (red). **b-c** Quantification of LVD (**b**) and LVA (**c**) (n=5/group). **d** Effects of VEGFC and VEGFD on tubule formation of primary LECs were detected by tubule formation experiment. **e** Immunofluorescence staining of VEGFC in CCl_4_- and BDL-induced fibrotic livers. **f-g** Quantification of mRNA and protein level of vegfc in CCl_4_-induced fibrotic liver by RT-qPCR (**f**) and western blot (**g**). **h** IHC staining of VEGFC in human fibrotic liver. **i** Serum VEGFC level between healthy control and fibrotic patients were assessed by ELISA (n=20-57/group). ns, not significant; *, P<0.05; ****, P<0.0001.

It’s reported that VEGFC was secreted by macrophages in ischemic liver ^[11]^. We first detected expression of VEGFC in macrophages by mIHC. The results showed that most VEGFC co-localized with macrophages in both human and murine fibrotic livers, indicating that VEGFC is primarily expressed by macrophages (Fig. 3a-b). Next, we depleted macrophages by injection of clodronate liposomes. As shown in Fig. 3c-e, depletion of macrophages resulted in significant down-regulation of VEGFC and inhibited lymphangiogenesis. Interestingly, we also found that T cells expressed a small part of VEGFC in human and murine fibrotic livers (Supplementary Fig. 4). Macrophages can be classified into M1 and M2 types. We wondered which type of macrophages predominantly secreted VEGFC. We identified M1 macrophages as the primary VEGFC producers instead of M2 macrophages in fibrotic livers by mIHC (Fig. 3F). Similarly, *in vitro* assays showed higher VEGFC expression in M1 than M2 and M0 (Fig. 3g-r).

**Fig. 3.**
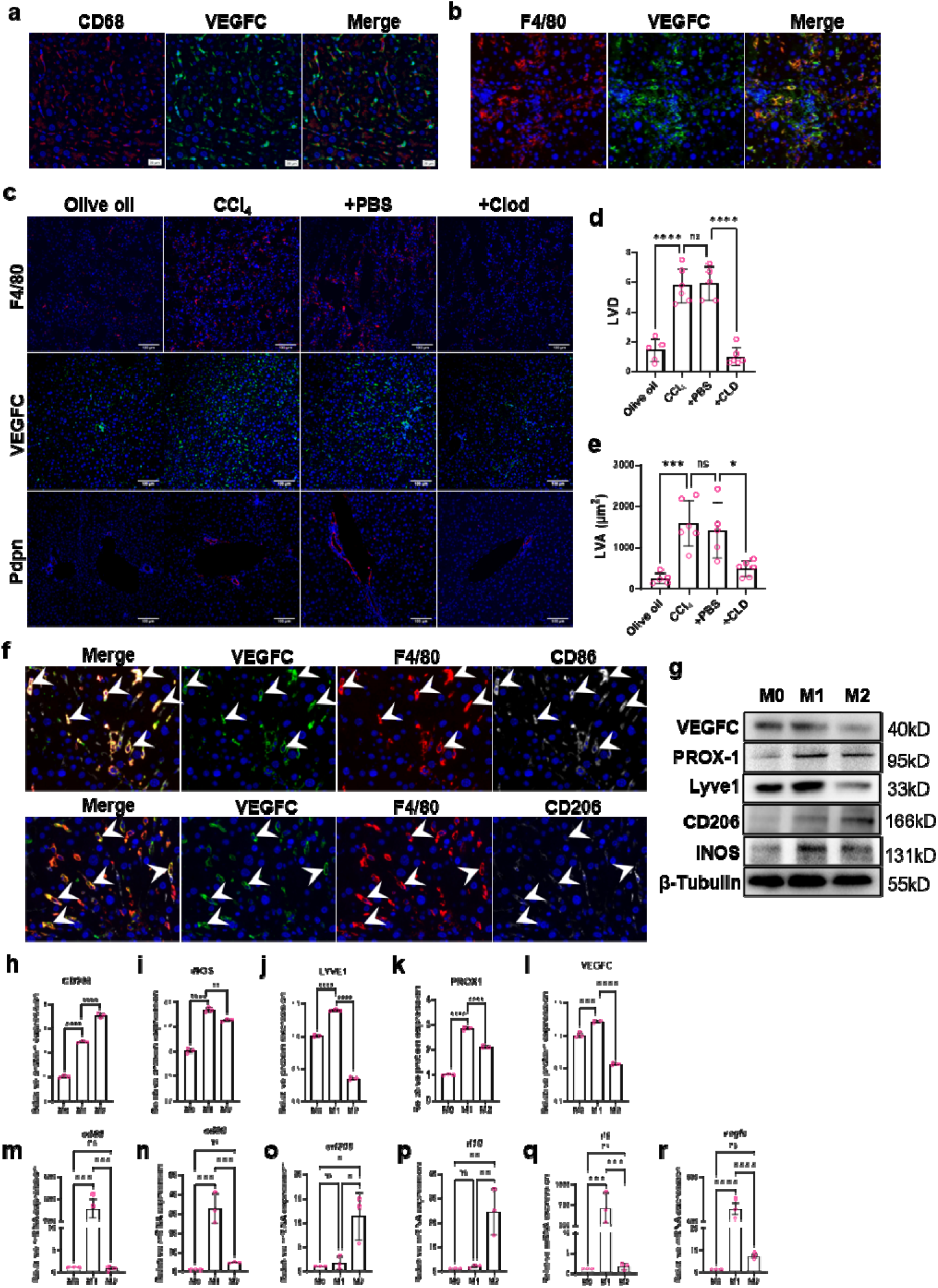
VEGFC mainly originates from M1 macrophages. **a** Co-localization of VEGFC (green) and macrophages indicated by CD68 (red) in human fibrotic liver was detected by mIHC. **b** Co-localization of VEGFC (green) and macrophages indicated by F4/80 (red) was detected by mIHC. **c-e** Macrophages were depleted by injection of Clodronate Liposomes (Clod). Immunofluorescence staining of F4/80, VEGFC and Pdpn (**c**) and quantification of LVD (**d**) and LVA (**e**). N=5-6/group. **f** Co-localization of VEGFC and M1 (marked by F4/80 and CD86) or M2 (marked by F4/80 and CD206) was detected by mIHC. **g-m** After THP-1 cells were induced by PMA into macrophages (M0), they were further polarized into M1 and M2 subtypes. Quantitative analysis of M1 markers (iNOS, CD80 and CD86), M2 markers (CD206, IL10 and IL6), VEGFC, and lymphatic markers (PROX1 and lyve1) was performed and analyzed using Western blot (**g-l**) and qRT-PCR (**m-r**). n=3/group. Ns, not significant; *, P<0.05; **, P<0.01; ***, P<0.001; ****, P<0.0001.

To further investigate the correlation between serum VEGFC levels and clinical characteristics of liver fibrosis, this study employed the median split method to divide 57 chronic liver fibrosis patients into high VEGFC (n=29) and low VEGFC (n=28) groups. Independent samples t-tests and chi-square tests revealed no statistically significant differences in baseline characteristics (e.g., age and gender distribution) between the two groups. The high-VEGFC group exhibited more pronounced features of hepatic injury, including significantly elevated AST (P<0.0001), ALT (P=0.005), and bile acid levels (P<0.0001). Concurrently, these patients demonstrated more evident lipid metabolism disorders, characterized by significantly increased LDL (P<0.0001) and a decreasing trend in HDL (P=0.079) despite lacking statistical significance. Notably, clinical staging analysis showed more advanced disease progression in the high-VEGFC group (P=0.049) (Table S2).

### 3.3 Lymphangiogenesis aggravates liver fibrosis

As the role of lymphangiogenesis remains controversial in liver fibrosis, we then verified its roles and importance in fibrogenesis. In addition to elevated serum ALT and AST, worse liver injury, increased ECM, more activated HSCs and elevated inflammatory cytokines were detected in VEGFC-treated group (Fig. 4). We also detected inhibited liver regeneration as indicated by reduction of proliferating hepatocytes (Fig. 4a, e). These results suggested that increased lymphangiogenesis by VEGFC injection aggravates liver fibrosis. It’s reported that angiogenesis plays key roles in liver fibrosis ^[12]^. We asked if VEGFC aggravates liver fibrosis by affecting angiogenesis. Staining of blood vessels and hepatic sinusoids confirmed little changes in angiogenesis after VEGFC treatment (Fig. 4a, d). Similar results were observed in BDL-induced liver fibrosis (Supplementary Figure 5). Moreover, VEGFC injection during fibrosis regression due to CCl_4_ cessation reversed fibrosis resolution, further confirming importance of lymphangiogenesis in liver fibrogenesis (Supplementary Fig. 6).

**Fig. 4.**
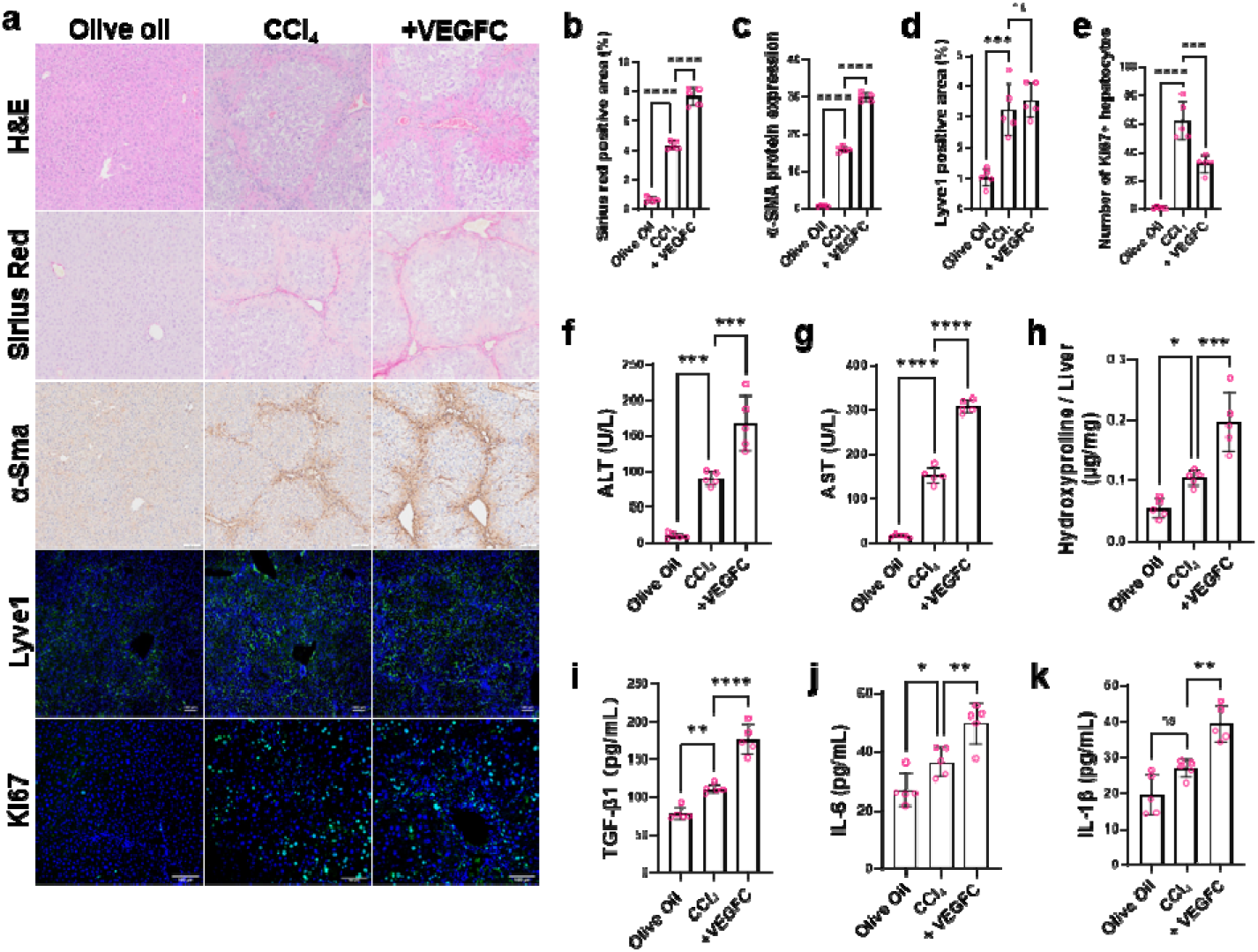
VEGFC-mediated lymphangiogenesis aggravated liver fibrosis. **a-e** Pathological section examination, IHC staining of α-sma and immunofluorescence staining of Lyve1 and Ki67 and the corresponding quantitative analysis. **f-g** ALT and AST were assessed. H. Hydroxyproline was assessed and adjusted to liver weight. **i-k** Tissue content of inflammatory cytokines including TGF-β1, IL-6 and IL-1β were quantified by Elisa. N=5/group. Ns, not significant; *, P<0.05; **, P<0.01; ***, P<0.001; ****, P<0.0001.

VEGFR3 is the receptor of VEGFC. We detected increased VEGFR3 expression in lymphatic vessels in fibrotic livers (Supplementary Fig. 7). Hence, we inhibited lymphangiogenesis using MAZ-51, a VEGFR3 inhibitor. Reduced lymphangiogenesis was confirmed by decreased LVA and LVD, and alleviated liver fibrosis were observed in MAZ-51-treated group, as indicated by reduced ECM content and inflammatory factors and restored liver functions (Fig. 5). Moreover, MAZ-51 abolished VEGFC-induced lymphangiogenesis and subsequent fibrosis, further confirming the importance of lymphangiogenesis in fibrosis (Fig. 6). These results suggested vital roles of lymphangiogenesis in hepatic fibrogenesis.

**Fig. 5.**
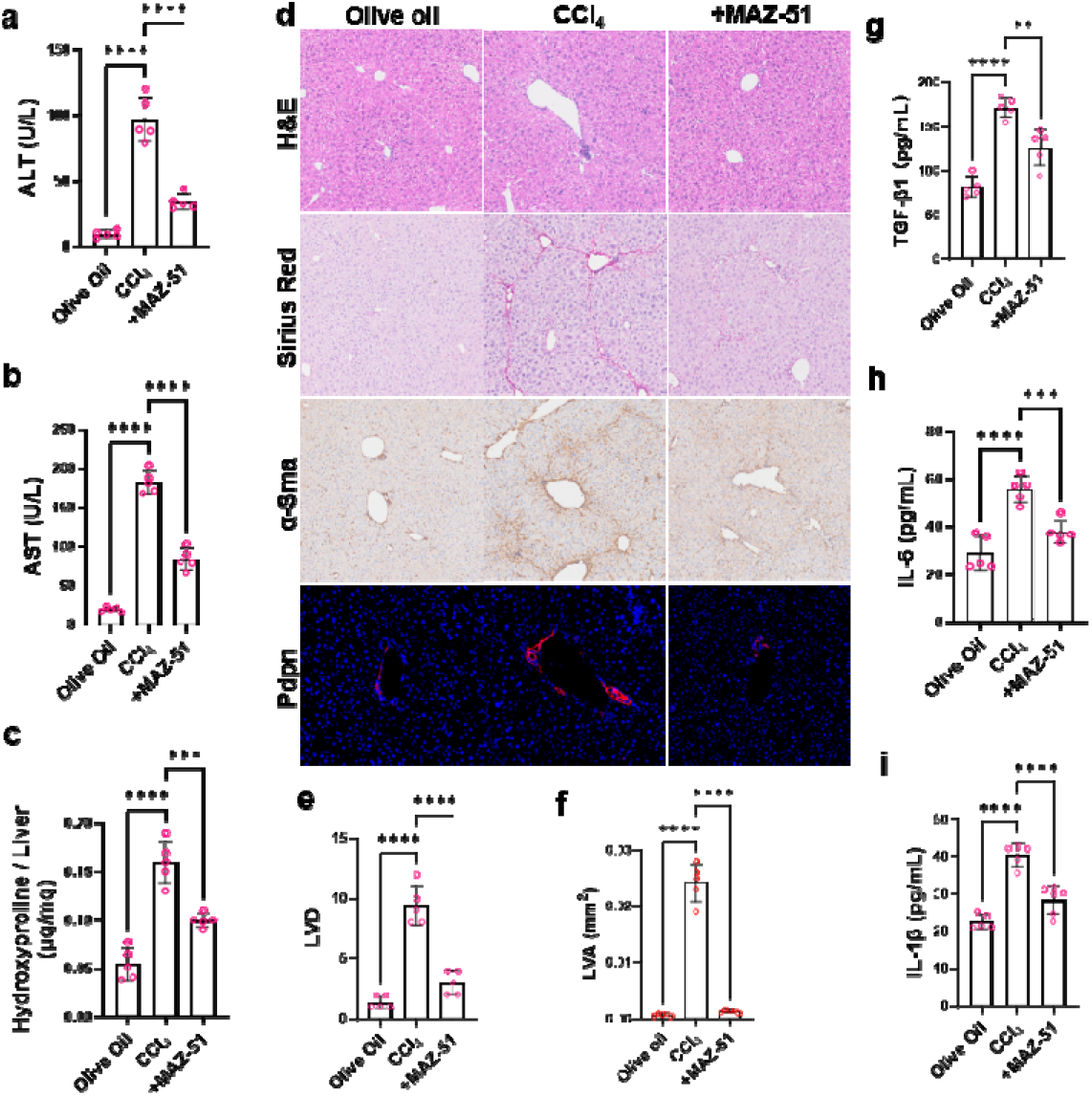
Decreased lymphangiogenesis and relieved fibrosis by MAZ-51 treatment. **a-b** ALT (**a**) and AST (**b**) were assessed. **c** Hydroxyproline was assessed and adjusted to liver weight. **d** Pathological section examination and IHC staining of were applied to assess liver injury and fibrosis. Lymphatic vessels were marked by immunofluorescence staining of pdpn. **e-f** Quantitative analysis of LVD (**e**) and LVA (**f**). **g-i** Tissue content of inflammatory cytokines including TGF-β1 (**g**), IL-6 (**h**) and IL-1β (**I**) were quantified by Elisa. N=5-6/group. **, P<0.01; ***, P<0.001; ****, P<0.0001.

**Fig. 6.**
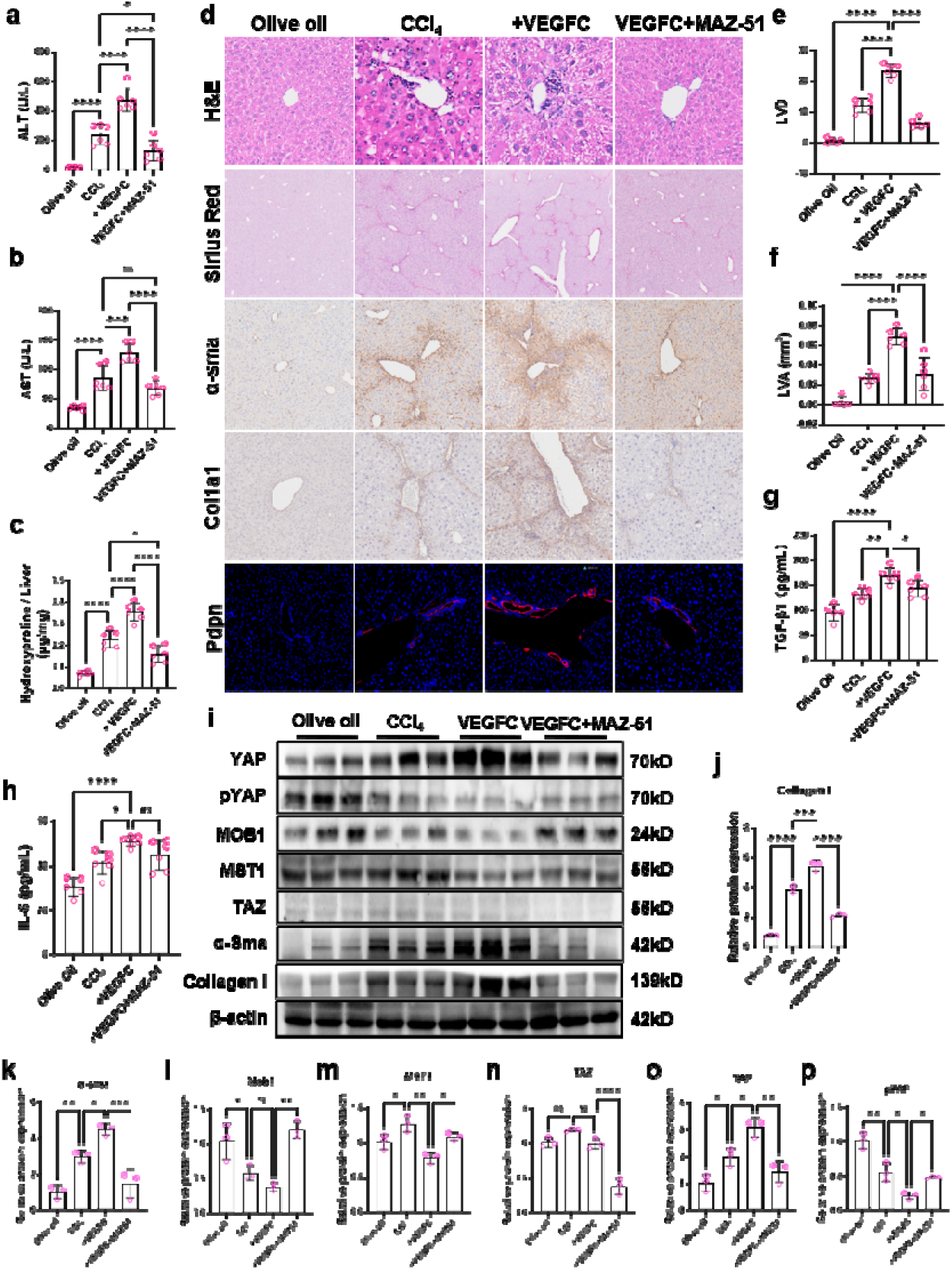
MAZ-51 reversed VEGFC-mediated fibrogenesis via modulating lymphangiogenesis. **a-b** ALT (**a**) and AST (**b**) were assessed. **c** Hydroxyproline was assessed and adjusted to liver weight. **d** Pathological section examination and IHC staining of α-sma and col1a1 were applied to assess liver injury and fibrosis. Lymphatic vessels were marked by immunofluorescence staining of pdpn. **e-f** Quantitative analysis of LVD (**e**) and LVA (**f**). **g-h** Tissue content of inflammatory cytokines including TGF-β1 (**g**) and IL-6 (**h**) were quantified by Elisa. **i-p** Hippo/YAP signaling pathway, α-sma and collagen I were measured by immunoblotting and normalized to β-actin. N=5-6/group. Ns, not significant; *, P<0.05; **, P<0.01; ***, P<0.001; ****, P<0.0001.

### 3.4 LECs-derived MDK promotes HSCs activation

HSCs activation is the key event driving liver fibrosis. We questioned if lymphangiogensis promoted fibrosis via HSCs activation. Firstly, we detected how LECs affected HSCs activation *in vitro*. We found that conditioned medium of LECs activated HSCs, and medium from VEGFC-treated LECs further promoted its activation, while MAZ-51 abolished VEGFC-treated LECs’ effects on HSCs activation (Supplementary Fig. 8). To reveal how lymphangiogenesis promotes HSCs activation, we conducted scRNA-seq analysis and found that interaction between LECs and HSCs was enhanced and MDK was specifically up-regulated when LECs acts on HSCs in cirrhotic patients (Fig. 7a). As the role of MDK in liver fibrosis is still unknown, we first verified its clinical significance by examining the plasma level and expression of MDK in livers of cirrhotic patients. The results showed that both plasma and liver expression of MDK was significantly increased in cirrhotic patients in contrast to its low expression in controls (Fig. 7b-d). More importantly, when patients were stratified into high- and low-expression groups based on median MDK levels for clinicopathological analysis (Table S3), the high-MDK group demonstrated significantly worse serological profiles, including Older age (P<0.001), markedly elevated liver injury markers, i.e. ALT (P=0.045), ALP (P=0.042), γ-GGT (P<0.0001), ADA (P=0.001), and TBIL (P=0.001), impaired synthetic function indicated by significantly reduced ALB (P=0.001) and TP (P=0.021), decreased HDL (P=0.037) and more advanced clinical staging (P<0.0001). These findings confirm the association between MDK and liver fibrosis severity.

**Fig. 7.**
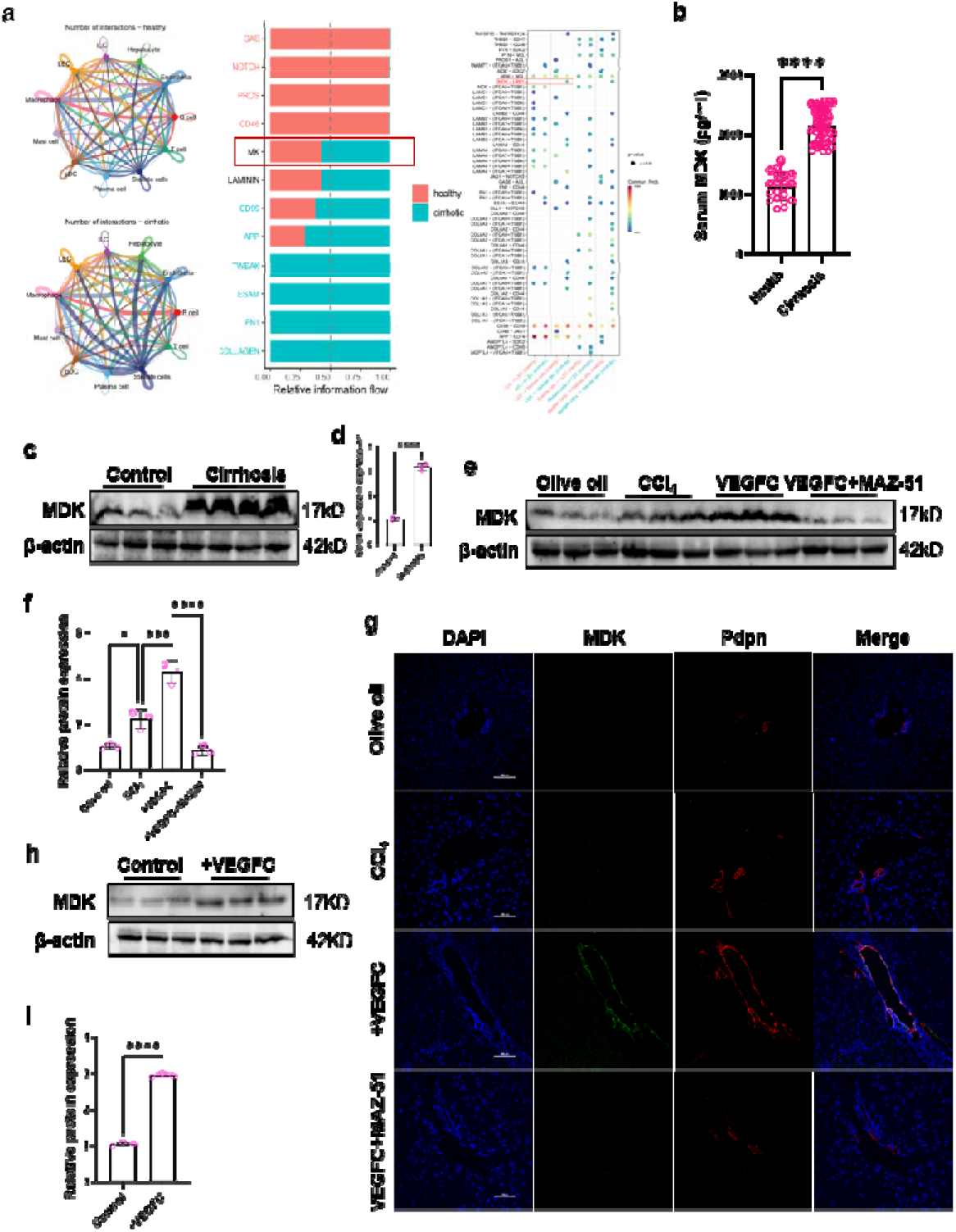
Increased MDK in LECs during liver fibrogenesis. **a** Analysis of the interaction between LECs and HSCs in cirrhosis based on the GSE136103 dataset. **b** Serum MDK level was assessed by Elisa (n=20-58/group). **c-f** Expression of VEGFC in human control and cirrhotic livers (**c-d**) and murine livers (**e-f**) was detected by immunoblotting. **g** Co-localization of MDK (green) and Pdpn (red) was detected by mIHC. The nuclei were stained by DAPI. **h-i** Primary LECs were treated with VEGFC, and the expression of MDK was detected by immunoblotting.*, P<0.05; ***, P<0.001; ****, P<0.0001.

Next, we found that MDK expression was significantly increased in murine fibrotic liver (Fig. 7e-f). Moreover, prompted lymphangiogenesis by VEGFC accompanied with increased MDK, while inhibited lymphangiogenesis by MAZ-51 resulted in reduced MDK (Fig. 7e-f). To elucidate specific expression of MDK in LECs, we examined co-localization of MDK and lymphatic vessels *in vivo* by mIHC (Fig 7g). Identically, *in vitro* study showed elevated MDK in VEGFC-treated LECs (Fig. 7h-i). These results suggested increased MDK expression in LECs during lymphangiogenesis.

Further, we examine the effects of MDK on HSCs activation. *In vitro* assay revealed that exogenous MDK was able to promote activation of HSCs (Fig. S9a). Identically, *in vivo* study showed that inhibition of MDK by its inhibitor-iMDK significantly reduced activated HSCs and ECM content, thus ameliorating liver fibrosis (Fig. S9b-d, g-j). Notably, we also observed inhibited lymphangiogenesis in iMDK group (Fig. S9e-f). To confirm if lymphangiogenesis facilitate liver fibrogenesis via secretion of MDK, iMDK was applied when lymphangiogenesis was induced by VEGFC injection. In line with our speculation, inhibition of MDK attenuated augmented fibrosis by VEGFC treatment (Fig. 8a-g).

**Fig. 8.**
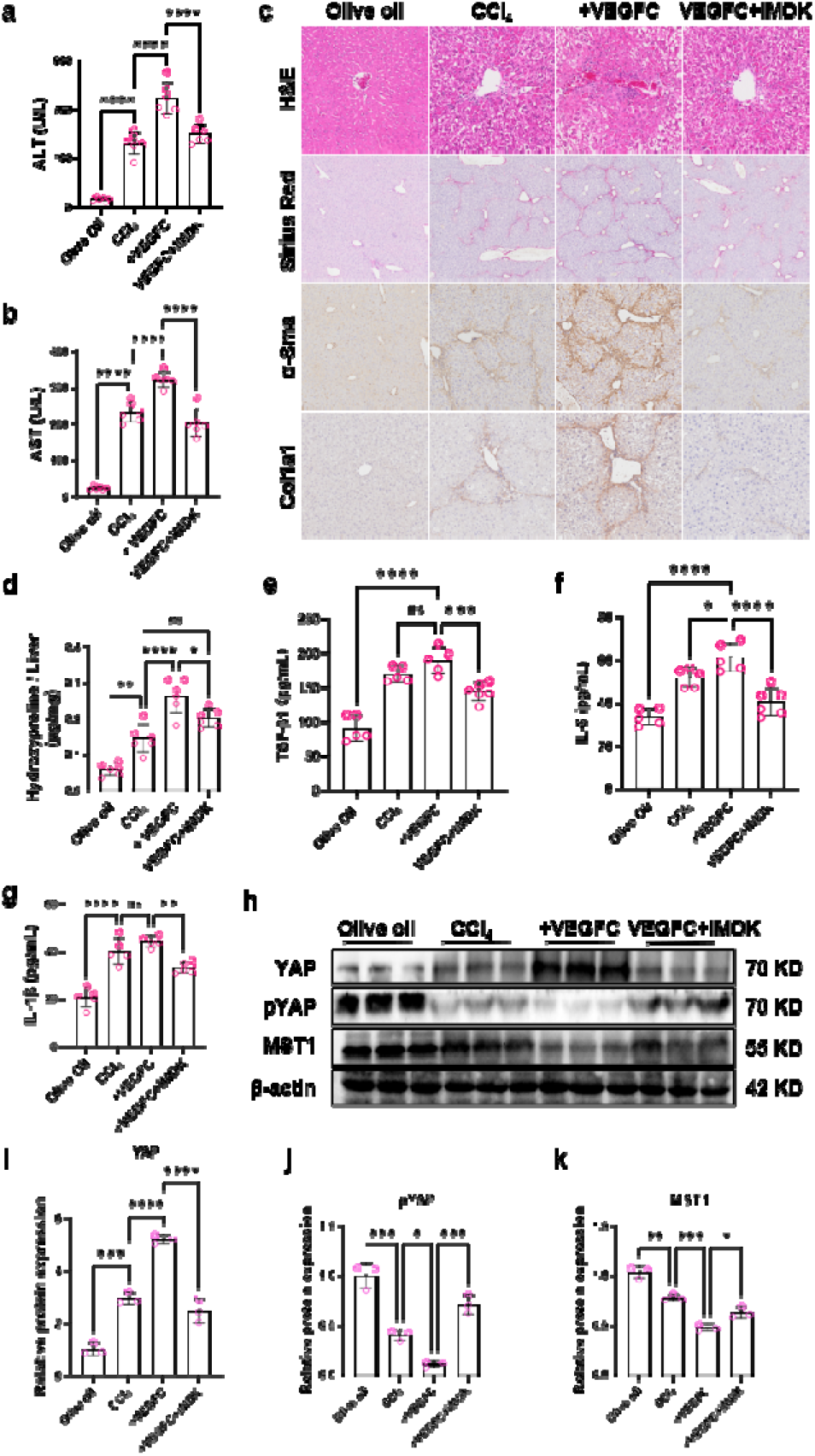
Inhibition of MDK reversed fibrosis induced by VEGFC. **a-b** ALT (**a**) and AST (**b**) were assessed. **c** Pathological section examination and IHC staining of α-sma and col1a1 were applied to assess liver injury and fibrosis. **d** Hydroxyproline was assessed and adjusted to liver weight. **e-g** Tissue content of inflammatory cytokines including TGF-β1 (**e**), IL-6 (**f**) and IL-1β (**g**) were quantified by Elisa. **h-k** Hippo/YAP signaling pathway were measured by immunoblotting. N=5-6/group. Ns, not significant; *, P<0.05; **, P<0.01; ***, P<0.001; ****, P<0.0001.

### 3.5 MDK activates HSCs via Hippo/YAP pathway

YAP is critical to HSCs activation ^[13]^. We detected activation of YAP in human fibrotic livers (Figure S10a-d). Moreover, our data indicated that MDK activated YAP through inhibiting Hippo pathway during HSCs activation (Fig. S9a). In accordance, inhibition of MDK activated Hippo signaling pathway, thus deactivating YAP (Fig. S9g, k-n). As shown in Figure 6I-P and 8H-K, VEGFC-induced lymphangiogenesis aggravated liver fibrosis, accompanying with increased MDK, YAP activation and Hippo inhibition. And both MAZ-51 and iMDK treatment abolished effects of VEGFC and deactivated downstream YAP via evoking Hippo pathway. Likewise, condition medium of VEGFC-treated LECs activated YAP via inhibition of Hippo pathway, which could be blocked by MAZ-51 (Fig. S8). Importantly, verteporfin, the inhibitor of YAP, inhibited HSCs activation induced by condition medium of LECs (Fig. S10e). These data suggested that LECs-derived MDK activates HSCs via Hippo/YAP pathway.

## 4 Discussion

Lymphangiogenesis has been conclusively demonstrated in hepatic fibrogenesis, yet its pathophysiological significance remains a subject of ongoing debate. Our study confirms enhanced lymphangiogenesis in both clinical specimens and murine fibrotic livers, while also demonstrating its reduction during fibrosis regression. Crucially, we show that lymphangiogenesis exacerbates liver fibrogenesis and that its inhibition attenuates fibrosis, identifying it as a promising therapeutic target. Mechanistically, we reveal that LECs secrete MDK to activate YAP signaling, thereby inducing HSCs activation and driving fibrogenesis. Furthermore, we establish that VEGFC - the key driver of lymphangiogenesis in hepatic fibrosis - is primarily derived from M1 macrophages.

Recently, emerging evidence has progressively elucidated the pivotal involvement of lymphangiogenesis in the pathogenesis of multiorgan fibrotic disorders. For example, in unilateral ureteral obstruction (UUO)-induced renal fibrosis models, lymphatic vessels recruit immune cells via the CCL21/CCR7 axis, exacerbating local inflammatory responses and accelerating fibrosis ^[14]^. In pulmonary and cardiac fibrosis, LECs can undergo endothelial-to-mesenchymal transition (EMT), differentiating into collagen-secreting myofibroblasts that directly contribute to extracellular matrix deposition ^[15]^. In peritoneal fibrosis, lymphatic dysfunction leads to impaired drainage capacity, thereby promoting fibrotic progression ^[16]^. Interestingly, however, some studies have reported protective effects of lymphangiogenesis. In specific models of cardiac ^[17]^, renal ^[18]^, and pulmonary fibrosis ^[19]^, lymphatic vessels may alleviate fibrosis by draining interstitial fluid, clearing immune cells, and reducing tissue edema. We speculate that the role of lymphangiogenesis in fibrotic progression may be highly dependent on organ-specific microenvironment and disease stages.

Studies report cell-type-specific sources of VEGFC that vary across liver diseases. In the liver ischemia-reperfusion model, macrophages are the primary VEGFC producers ^[11]^, whereas Schwann cells - not macrophages - predominantly express VEGFC in a rat partial portal vein ligation (PPVL) model ^[5]^. In a mouse NASH model, VEGFC is primarily secreted by LECs and PECs (portal endothelial cells). However, since sequencing was performed exclusively on endothelial cells, the possibility of VEGFC secretion by non-endothelial cell populations cannot be excluded ^[8]^. Here, we demonstrate that M1 macrophages are the dominant VEGFC-expressing population in liver fibrosis. Notably, we also detected VEGFC in T cells, suggesting their potential involvement in lymphangiogenesis regulation in liver fibrosis. Interestingly, the origin of VEGFC varies across different organs. Lin et al. Reported that mRNA expression of VEGFC and VEGFR3 was found most significantly up-regulated in cardiac LECs ^[20]^. Lim et al. found that platelets stimulate lymphangiogenesis via VEGFC release after wounding ^[21]^. Moreover, VEGFC is not even the factor mediating lymphangiogenesis in certain disease contexts. For instance, previous studies have demonstrated that tumor-associated neutrophils promote lymphangiogenesis through VEGFA and MMP9 secretion, rather than via VEGFC or VEGFD, in bladder cancer ^[22]^. These findings suggest distinct upstream regulatory mechanisms of lymphangiogenesis across different disease contexts.

Although we have demonstrated that the primary source of VEGFC and LECs activates HSCs through the MDK/YAP signaling pathway during lymphangiogenesis, the precise mechanism by which VEGFC promotes lymphangiogenesis remains to be elucidated. In renal fibrosis, VEGFC promotes lymphangiogenesis via mediating macrophages-LECs transition, which involves autophagy in macrophages ^[23]^. It’s also reported that autophagy is essential for LECs to response to VEGFC and lymphangiogenesis in a corneal wound healing model ^[24]^. Koltowska et al. reported VEGFC-SoxF-Mafba signaling pathway for embryonic lymphangiogenesis in zebrafish ^[25]^. Overall, there is limited research on how the VEGFC/VEGFR3 axis promotes lymphangiogenesis, and the mechanisms remain unclear. New evidence is needed to elucidate the relevant mechanisms.

Macrophages participate in multiple signaling pathways during liver fibrogenesis, playing pivotal roles in both the initiation and progression of fibrosis, while exhibiting remarkable functional complexity and diversity. In this study, we demonstrated that M1-type macrophages promote lymphangiogenesis through the secretion of VEGFC. Zhang Y. et al. reported that VEGFC/VEGFR3 signaling drives the transdifferentiation of M1 macrophages into LECs in renal fibrosis ^[23]^. Similarly, we observed colocalization of macrophages and LECs in human fibrotic liver (data not shown), suggesting the potential for macrophage-to-LEC transition in liver fibrosis.

MDK is a secreted growth factor involved in fundamental cellular processes including proliferation and migration. It exhibits high expression during embryogenesis, gradually declines during early postnatal development, and becomes nearly undetectable in healthy adults ^[26]^. MDK has been identified as a potential serum biomarker for hepatocellular carcinoma (HCC), demonstrating superior sensitivity to α-fetoprotein (AFP) in early-stage HCC diagnosis ^[27]^. This study revealed that serum MDK levels were closely associated with liver fibrosis progression, suggesting MDK as a potential molecular biomarker for evaluating fibrosis advancement.

Current research on hepatic MDK primarily focuses on its applications in HCC, either as a diagnostic biomarker or therapeutic target ^[28]^. In other liver disease models, limited studies have reported MDK involvement. Cadmium intoxication was shown to upregulate hepatic MDK expression, though the underlying mechanisms remain unexplored ^[29]^. Using rat primary liver sinusoidal endothelial cells (LSECs), Wu L et al. Found that MDK promotes capillarization through the integrin α4/NF-κB pathway ^[30]^. Regarding liver fibrosis, a single study reported that quercetin could suppress MDK expression in activated HSCs and induce their apoptosis ^[31]^, suggesting a potential role of MDK in fibrogenesis. Our data indicate that LECs derived-MDK activates HSCs via Hippo/YAP signal pathway, thus promoting fibrosis.

## 5 Conclusions

In summary, our study establishes that M1 macrophages promote lymphangiogenesis through VEGFC secretion during liver fibrogenesis. Lymphangiogenesis exacerbate fibrosis by producing MDK, which activates HSCs via the Hippo/YAP signaling pathway. Crucially, pharmacological inhibition of lymphangiogenesis through VEGFC/VEGFR3 blockade significantly attenuates fibrosis, highlighting both its pathophysiological importance and therapeutic potential in liver fibrosis.

## Statements

### Data availability statement

The data which support the findings of this study are available from the corresponding author upon reasonable request.

### Ethics statement

The animal study was approved by the Committee on Animal Research of Sichuan University. The study was conducted in accordance with the local legislation and institutional requirements.

### Author information

Dan Wang and Dan Long contributed equally to this work.

### Author Contributions

**Dan Wang:** Investigation, Data curation, Methodology, Software. **Dan Long:** Investigation, Methodology. **Ying Zhao:** Investigation, Data curation. **Dian Li:** Investigation. **Fenglin Xiong**: Data curation. **Ziwei Huang:** Data curation. **Liping Yang:** Data curation. **Qing Zheng:** Investigation. **Yanni Zhou:** Investigation, Data curation, Methodology, Funding acquisition, Writing - original draft. **Yonghua Chen & Li Feng:** Conceptualization, Funding acquisition, Project administration, Supervision, Writing - review & editing.

### Funding

This study was supported by the grant from the National Natural Science Foundation of China (No. 82100649) and the Science and Technology Bureau of Chengdu (2024-YF05-00486-SN).

### Conflict of interests

The authors have no relevant financial or non-financial interests to disclose.

## Abbreviations

CCl_4_: carbon tetrachloride
TAA: Thioacetamide
Pdpn: podoplain
MDK: Midkine
HSCs: hepatic stellate cells
ECM: extracellular matrix
HCC: hepatocellular carcinoma
LVD: lymphatic vessel density
PDGF-D: platelet-derived growth factor-D
ACR: acute cellular rejection
BDL: bile duct ligation
LECs: lymphatic endothelial cells
NASH: non-alcoholic steatohepatitis
scRNA-seq: single-cell RNA sequencing
ALT: alanine transaminase
AST: aspartate aminotransferase
PMA: Phorbol myristate acetate
LVA: lymphatic vessel area
mIHC: multiplexing immunohistochemistry
DAB: 3,3⍰-diaminobenzidine
AST: aspartate aminotransferase
ALT: alanine aminotransferas
γ-GGT: γ-gamma-glutamyl transferase
ALP: alkaline phosphatase
BAs: bile acids
ADA: adenosine deaminase
TBIL: total bilirubin
TP: total protein
ALB: albumin
TT: thrombin time
APTT: activated partial thromboplastin time
PT: prothrombin time
D-D: D-dimer
FIB: fibrinogen
HDL: high density lipoprotein
LDL: low density lipoprotein
UUO: unilateral ureteral obstruction
EMT: endothelial-to-mesenchymal transition
PPVL: partial portal vein ligation
PECs: portal endothelial cells
AFP: α-fetoprotein
LSECs: liver sinusoidal endothelial cells.

## Notes

### Competing Interest Statement

The authors have declared no competing interest.

## References

1. Ohtani O, Ohtani Y. Lymph circulation in the liver. Anatomical record (Hoboken, NJ : 2007) 2008; 291(6):643–652.

2. Cadamuro M, Brivio S, Mertens J, Vismara M, Moncsek A, Milani C, et al. Platelet-derived growth factor-D enables liver myofibroblasts to promote tumor lymphangiogenesis in cholangiocarcinoma. Journal of hepatology 2019; 70(4):700–709.

3. Sakamoto K, Ogawa K, Tamura K, Honjo M, Funamizu N, Takada Y. Prognostic Role of the Intrahepatic Lymphatic System in Liver Cancer. Cancers 2023; 15(7).

4. Ishii E, Shimizu A, Kuwahara N, Arai T, Kataoka M, Wakamatsu K, et al. Lymphangiogenesis associated with acute cellular rejection in rat liver transplantation. Transplantation proceedings 2010; 42(10):4282–4285.

5. Tanaka M, Jeong J, Thomas C, Zhang X, Zhang P, Saruwatari J, et al. The Sympathetic Nervous System Promotes Hepatic Lymphangiogenesis, which Is Protective Against Liver Fibrosis. The American journal of pathology 2023; 193(12):2182–2202.

6. Jeong J, Tanaka M, Yang Y, Arefyev N, DiRito J, Tietjen G, et al. An optimized visualization and quantitative protocol for in-depth evaluation of lymphatic vessel architecture in the liver. American journal of physiology Gastrointestinal and liver physiology 2023; 325(5):G379–g390.

7. Su T, Yang Y, Lai S, Jeong J, Jung Y, McConnell M, et al. Single-Cell Transcriptomics Reveals Zone-Specific Alterations of Liver Sinusoidal Endothelial Cells in Cirrhosis. Cellular and molecular gastroenterology and hepatology 2021; 11(4):1139–1161.

8. Burchill MA, Finlon JM, Goldberg AR, Gillen AE, Dahms PA, McMahan RH, et al. Oxidized Low-Density Lipoprotein Drives Dysfunction of the Liver Lymphatic System. Cellular and molecular gastroenterology and hepatology 2021; 11(2):573–595.

9. Huang S, Li B, Liu Z, Xu M, Lin D, Hu J, et al. Three-dimensional mapping of hepatic lymphatic vessels and transcriptome profiling of lymphatic endothelial cells in healthy and diseased livers. Theranostics 2023; 13(2):639–658.

10. Tamburini BAJ, Finlon JM, Gillen AE, Kriss MS, Riemondy KA, Fu R, et al. Chronic Liver Disease in Humans Causes Expansion and Differentiation of Liver Lymphatic Endothelial Cells. Frontiers in immunology 2019; 10:1036.

11. Nakamoto S, Ito Y, Nishizawa N, Goto T, Kojo K, Kumamoto Y, et al. Lymphangiogenesis and accumulation of reparative macrophages contribute to liver repair after hepatic ischemiareperfusion injury. Angiogenesis 2020; 23(3):395–410.

12. Li H. Angiogenesis in the progression from liver fibrosis to cirrhosis and hepatocelluar carcinoma. Expert review of gastroenterology & hepatology 2021; 15(3):217–233.

13. Du K, Hyun J, Premont RT, Choi SS, Michelotti GA, Swiderska-Syn M, et al. Hedgehog-YAP Signaling Pathway Regulates Glutaminolysis to Control Activation of Hepatic Stellate Cells. Gastroenterology 2018; 154(5):1465-1479.e1413.

14. Liu Z, Zhang C, Hao J, Chen G, Liu L, Xiong Y, et al. Eplerenone ameliorates lung fibrosis in unilateral ureteral obstruction rats by inhibiting lymphangiogenesis. Experimental and therapeutic medicine 2022; 24(4):623.

15. Terabayashi T, Ito Y, Mizuno M, Suzuki Y, Kinashi H, Sakata F, et al. Vascular endothelial growth factor receptor-3 is a novel target to improve net ultrafiltration in methylglyoxal-induced peritoneal injury. Laboratory investigation; a journal of technical methods and pathology 2015; 95(9):1029–1043.

16. Wang C, Yue Y, Huang S, Wang K, Yang X, Chen J, et al. M2b macrophages stimulate lymphangiogenesis to reduce myocardial fibrosis after myocardial ischaemia/reperfusion injury. Pharmaceutical biology 2022; 60(1):384–393.

17. Hasegawa S, Nakano T, Torisu K, Tsuchimoto A, Eriguchi M, Haruyama N, et al. Vascular endothelial growth factor-C ameliorates renal interstitial fibrosis through lymphangiogenesis in mouse unilateral ureteral obstruction. Laboratory investigation; a journal of technical methods and pathology 2017; 97(12):1439–1452.

18. El-Chemaly S, Malide D, Zudaire E, Ikeda Y, Weinberg BA, Pacheco-Rodriguez G, et al. Abnormal lymphangiogenesis in idiopathic pulmonary fibrosis with insights into cellular and molecular mechanisms. Proceedings of the National Academy of Sciences of the United States of America 2009; 106(10):3958–3963.

19. Lee AS, Lee JE, Jung YJ, Kim DH, Kang KP, Lee S, et al. Vascular endothelial growth factor-C and -D are involved in lymphangiogenesis in mouse unilateral ureteral obstruction. Kidney international 2013; 83(1):50–62.

20. Lin QY, Zhang YL, Bai J, Liu JQ, Li HH. VEGF-C/VEGFR-3 axis protects against pressure-overload induced cardiac dysfunction through regulation of lymphangiogenesis. Clinical and translational medicine 2021; 11(3):e374.

21. Lim L, Bui H, Farrelly O, Yang J, Li L, Enis D, et al. Hemostasis stimulates lymphangiogenesis through release and activation of VEGFC. Blood 2019; 134(20):1764–1775.

22. Zhang Q, Liu S, Wang H, Xiao K, Lu J, Chen S, et al. ETV4 Mediated Tumor-Associated Neutrophil Infiltration Facilitates Lymphangiogenesis and Lymphatic Metastasis of Bladder Cancer. Advanced science (Weinheim, Baden-Württemberg, Germany) 2023; 10(11):e2205613.

23. Zhang Y, Zhang C, Li L, Liang X, Cheng P, Li Q, et al. Lymphangiogenesis in renal fibrosis arises from macrophages via VEGF-C/VEGFR3-dependent autophagy and polarization. Cell death & disease 2021; 12(1):109.

24. Meçe O, Houbaert D, Sassano ML, Durré T, Maes H, Schaaf M, et al. Lipid droplet degradation by autophagy connects mitochondria metabolism to Prox1-driven expression of lymphatic genes and lymphangiogenesis. Nature communications 2022; 13(1):2760.

25. Koltowska K, Paterson S, Bower NI, Baillie GJ, Lagendijk AK, Astin JW, et al. mafba is a downstream transcriptional effector of Vegfc signaling essential for embryonic lymphangiogenesis in zebrafish. Genes & development 2015; 29(15):1618–1630.

26. Filippou PS, Karagiannis GS, Constantinidou A. Midkine (MDK) growth factor: a key player in cancer progression and a promising therapeutic target. Oncogene 2020; 39(10):2040–2054.

27. Lu Q, Li J, Cao H, Lv C, Wang X, Cao S. Comparison of diagnostic accuracy of Midkine and AFP for detecting hepatocellular carcinoma: a systematic review and meta-analysis. Bioscience reports 2020; 40(3).

28. Mamdouh S, Khorshed FEM, Hammad G, Magdy M, Abdelraouf A, Hemida E, et al. RNA Interference based Midkine Gene Therapy for Hepatocellular Carcinoma. Asian Pacific journal of cancer prevention :APJCP 2024; 25(7):2371–2379.

29. Yazihan N, Kocak MK, Akcil E, Erdem O, Sayal A. Role of midkine in cadmium-induced liver, heart and kidney damage. Human & experimental toxicology 2011; 30(5):391–397.

30. Wu L, Chen H, Eu C, Xing M, Fang H, Yang F, et al. Midkine mediates dysfunction of liver sinusoidal endothelial cells through integrin a4 and a6. Vascular pharmacology 2022; 147:107113.

31. Wu LC, Lu IW, Chung CF, Wu HY, Liu YT. Antiproliferative mechanisms of quercetin in rat activated hepatic stellate cells. Food & function 2011; 2(3–4):204–212.

