## Supplementary files for "M1 macrophage-mediated lymphangiogenesis aggravates liver fibrosis via MDK/YAP signaling pathway"

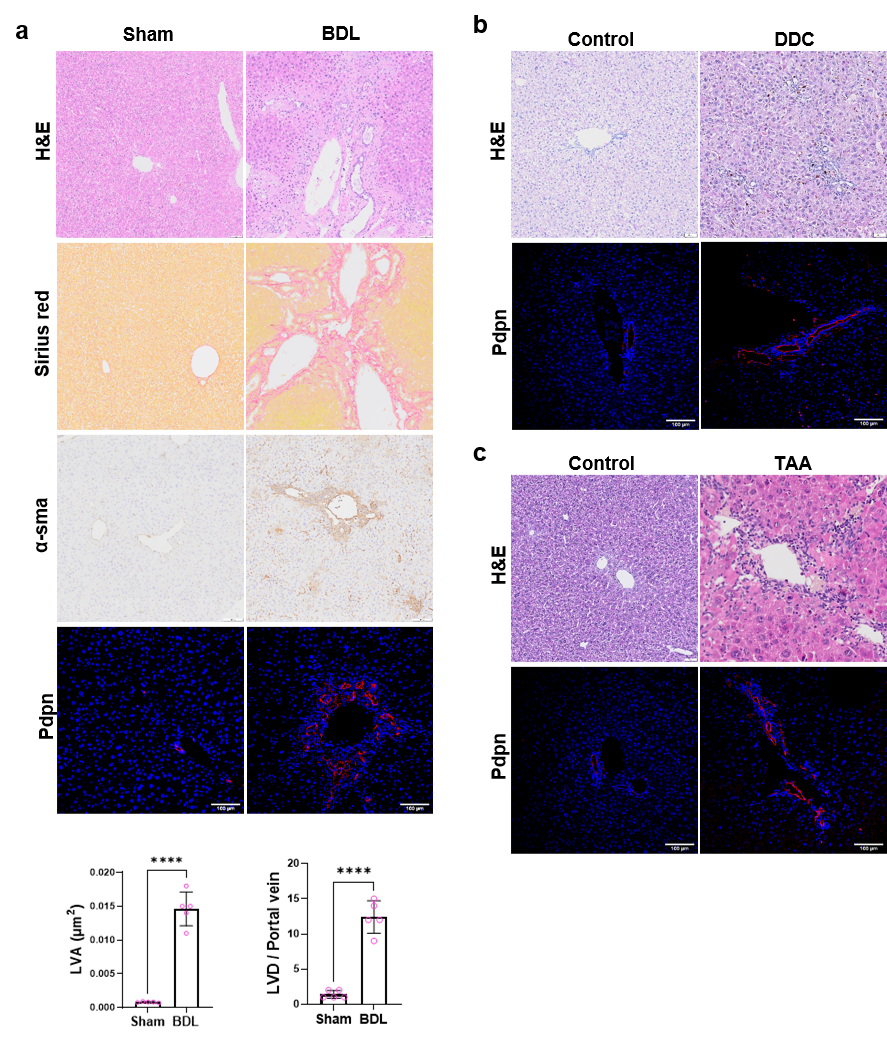

**Fig. S1** Increased lymphangiogenesis in fibrotic livers of various mouse models. **a-c** Pathological section examination and IHC staining of α-sma was conducted to confirm liver fibrosis induced by BDL (**a**), DDC (**b**) and TAA (**c**). Lymphatic vessels were marked by immunofluorescence staining of Pdpn.

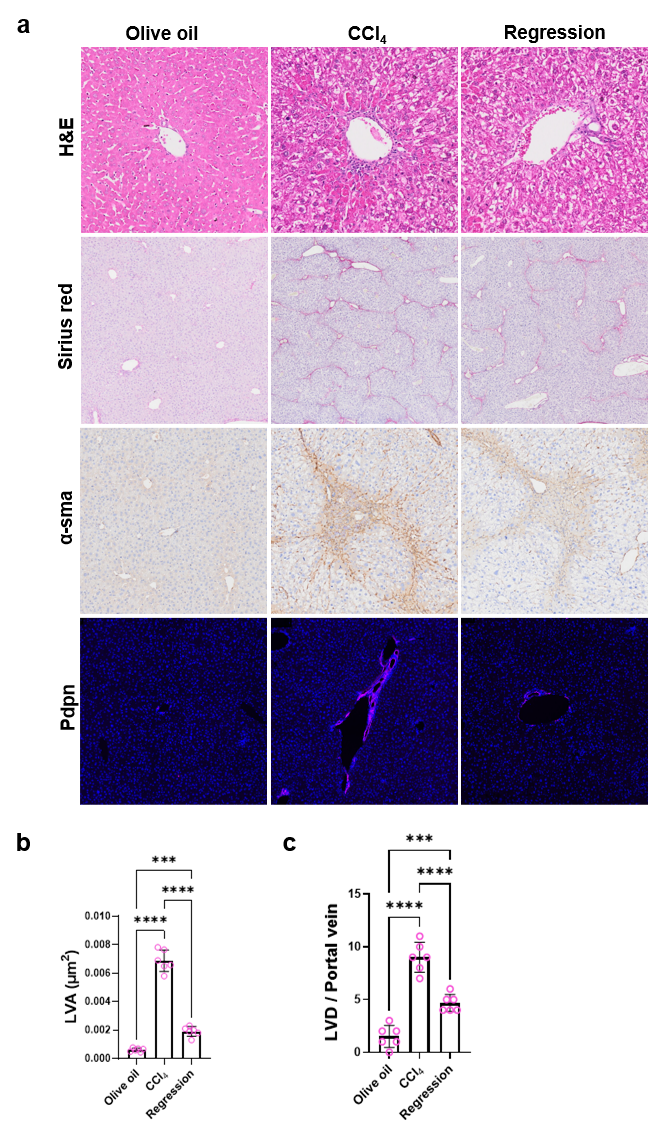

**Fig. S2** Reduced lymphangiogenesis during liver fibrosis regression. **a** Pathological section examination and IHC staining of α-sma was conducted to confirm liver fibrosis regression after cessation of CCl4. Lymphatic vessels were marked by immunofluorescence staining of Pdpn. **b-c** Quantitative analysis of LVA (**b**) and LVD (**c**). ***, P<0.001; ****, P<0.0001.

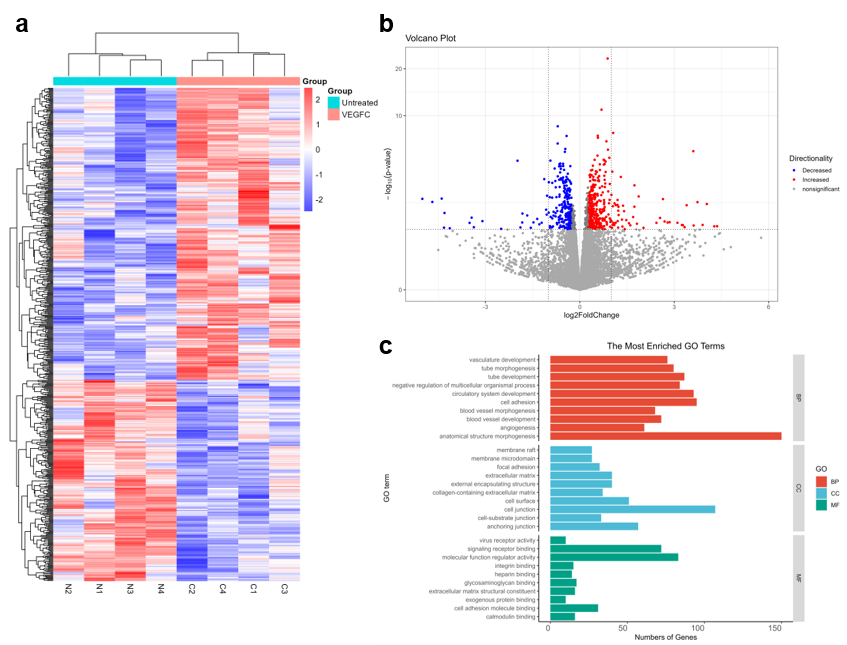

**Fig. S3** Transcriptomic analysis of effects on LECs by VEGFC treatment.

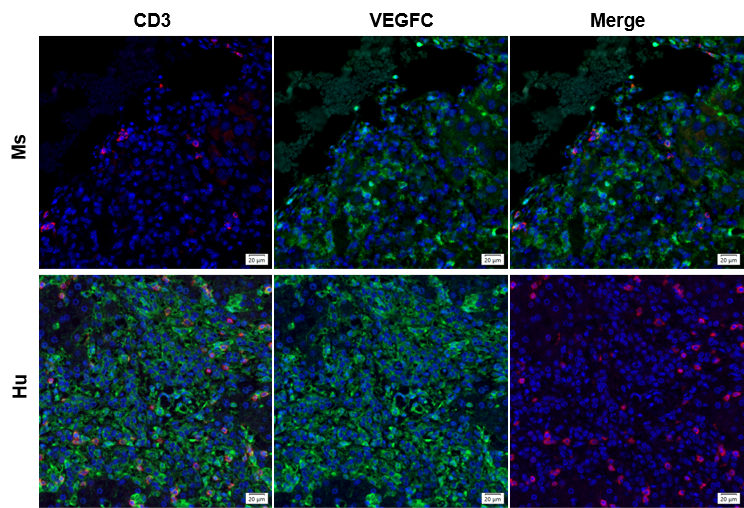

**Fig. S4** Co-localization of VEGFC and T cells in human cirrhotic liver by mIHC.

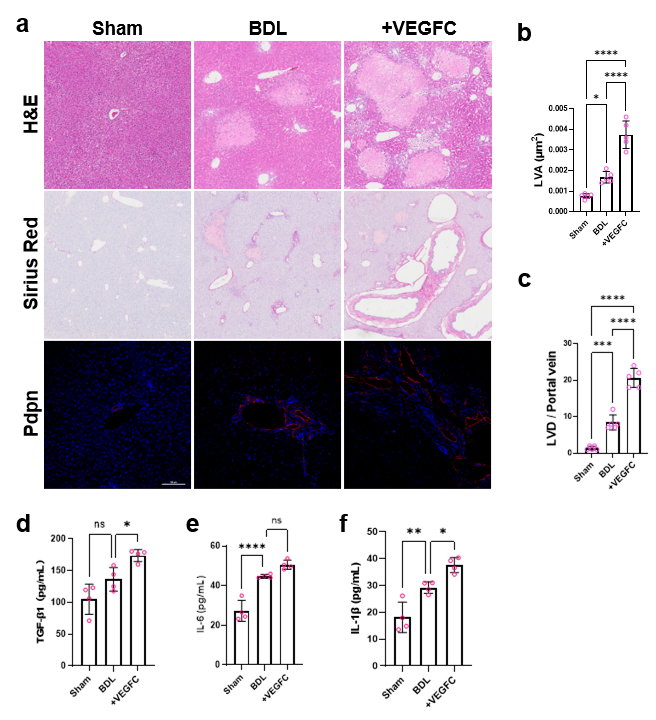

**Fig. S5** lymphangiogenesis promoted liver fibrosis in BDL-induced liver fibrosis. **a** Pathological section examination was applied to assess liver injury and fibrosis. **b-c** Quantitative analysis of LVA (**b**) and LVD (**c**). **d-f** Tissue content of inflammatory cytokines including TGF-β1 (**d**), IL-6 (**e**) and IL-1β (**f**) were quantified by Elisa. N=4/group. Ns, not significant; *, P<0.05; **, P<0.01; ***, P<0.001; ****, P<0.0001.

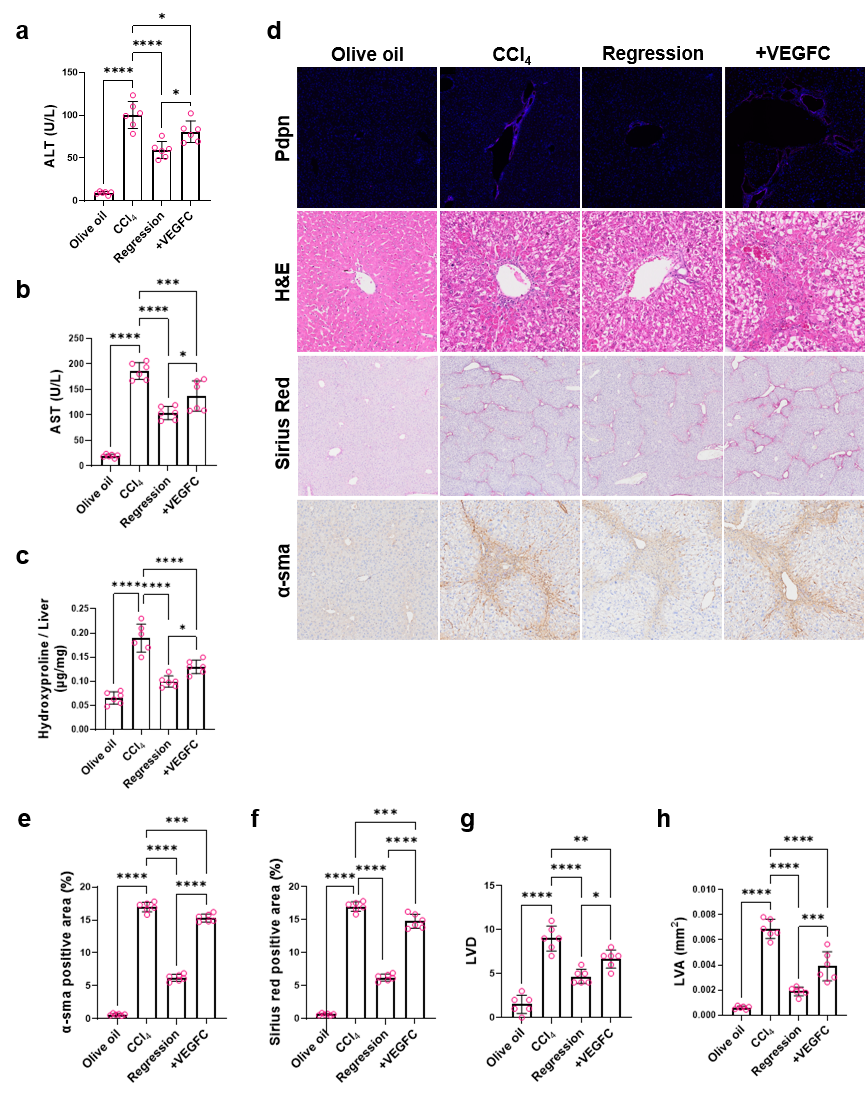

**Fig. S6** VEGFC-induced lymphangiogenesis reversed liver fibrosis regression. **a-b** ALT (**a**) and AST (**b**) were assessed. **c** Hydroxyproline was assessed and adjusted to liver weight. **d** Pathological section examination and IHC staining of α-sma and col1a1 were applied to assess liver injury and fibrosis. Lymphatic vessels were marked by immunofluorescence staining of Pdpn. **e-f** Quantification of α-sma (**e**) and sirius red (**f**) positive area. **g-h** Quantitative analysis of LVD (**g**) and LVA (**h**). N=6/group. *, P<0.05; **, P<0.01; ***, P<0.001; ****, P<0.0001.

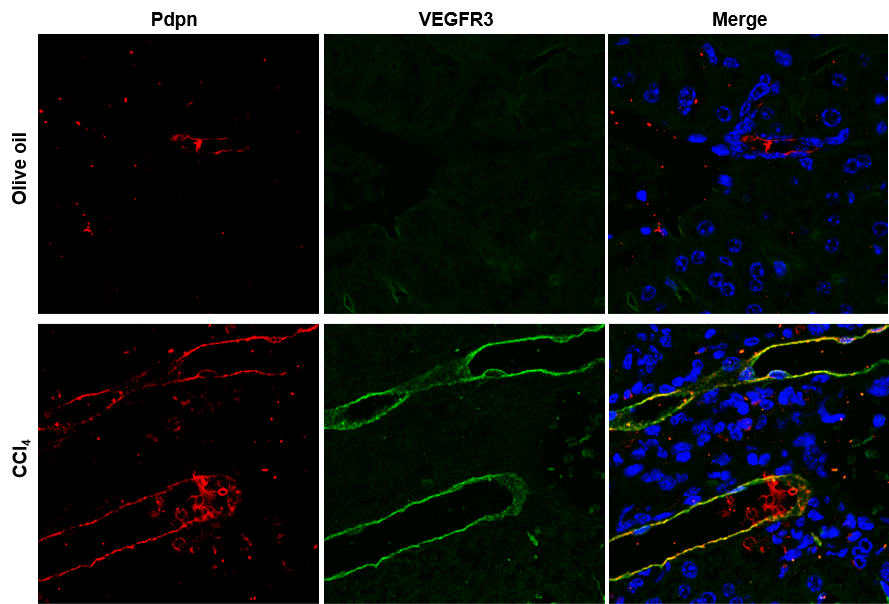

**Fig. S7** Expression of VEGFR3 in lymphatic vessels. Lymphatic vessels were marked by Pdpn (red). Co-localization of VEGFR3 and Pdpn was yellow.

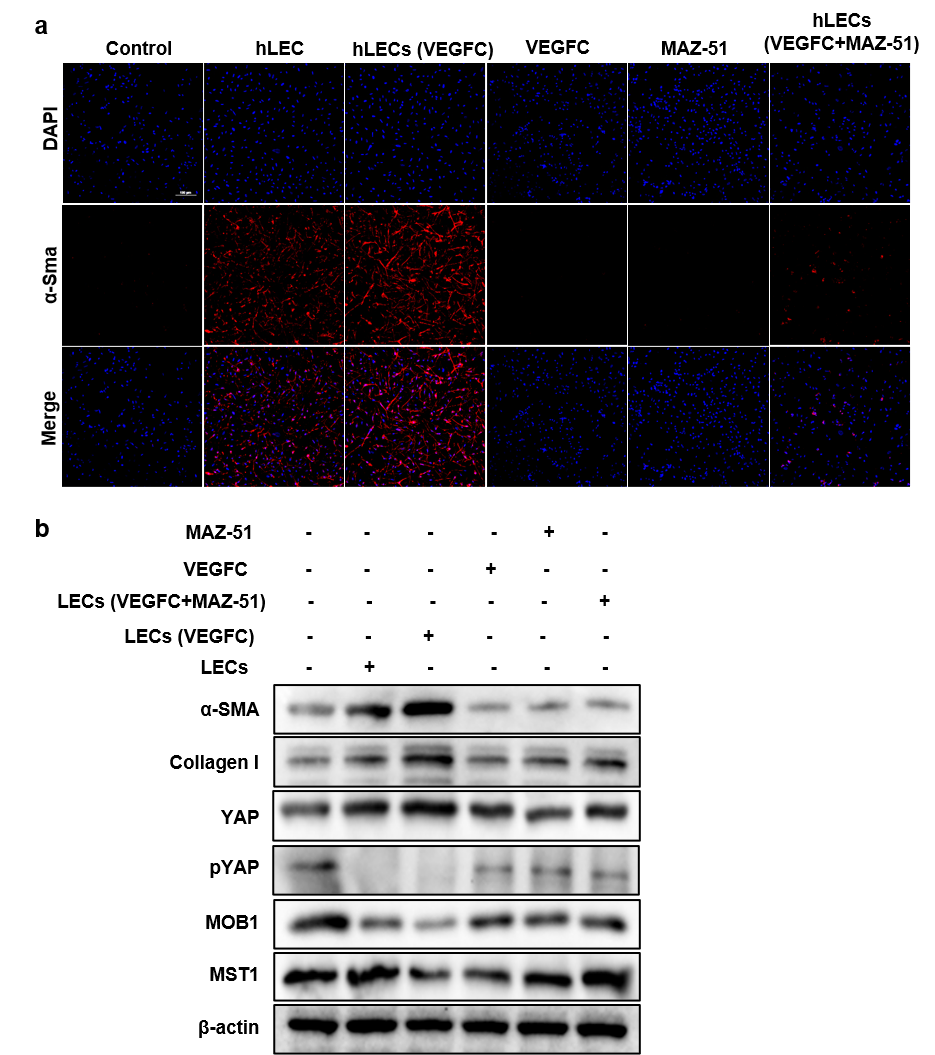

**Fig. S8** VEGFC-treated LECs activated HSCs. **a** Immunofluorescence staining of α-sma in each group. **b** Hippo/YAP signaling pathway, collagen I and α-sma were assessed by immunoblotting.

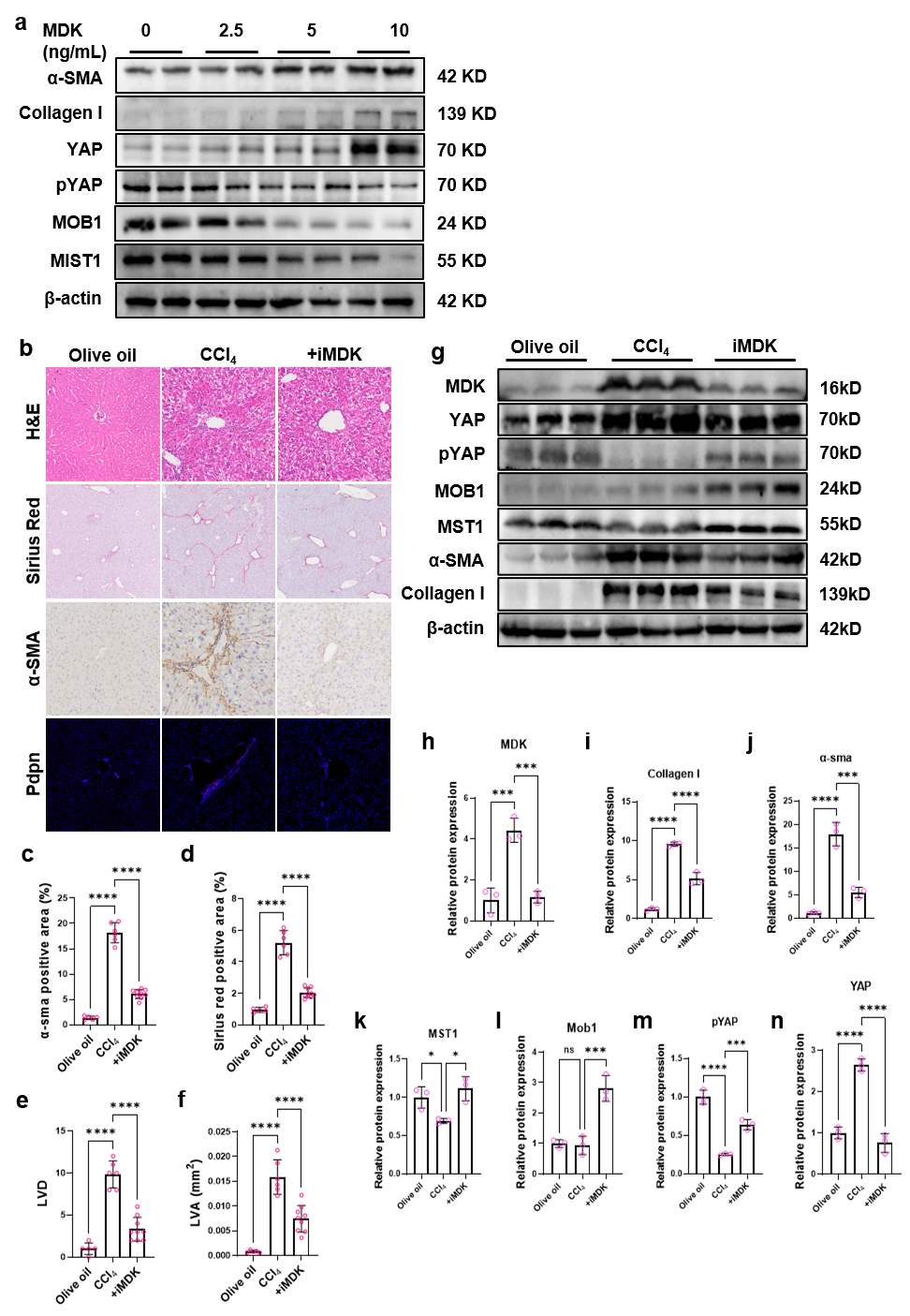

**Fig. S9** MDK activated HSCs. **a** MDK at different final concentrations was added to the culture medium of LX-2. Activation and Hippo/YAP signaling pathway of LX-2 was measured by immunoblotting. **b** Pathological section examination and IHC staining of α-sma and col1a1 were applied to assess liver injury and fibrosis. Lymphatic vessels were marked by immunofluorescence staining of Pdpn. **c-d** Quantification of α-sma (**c**) and sirius red (**d**) positive area. **e-f** Quantitative analysis of LVD (**e**) and LVA (**f**). **g-n** Hippo/YAP signaling pathway, MDK, collagen I and α-sma were assessed by immunoblotting and normalized to β-actin. Ns, not significant. *, P<0.05; ***, P<0.001; ****, P<0.0001.

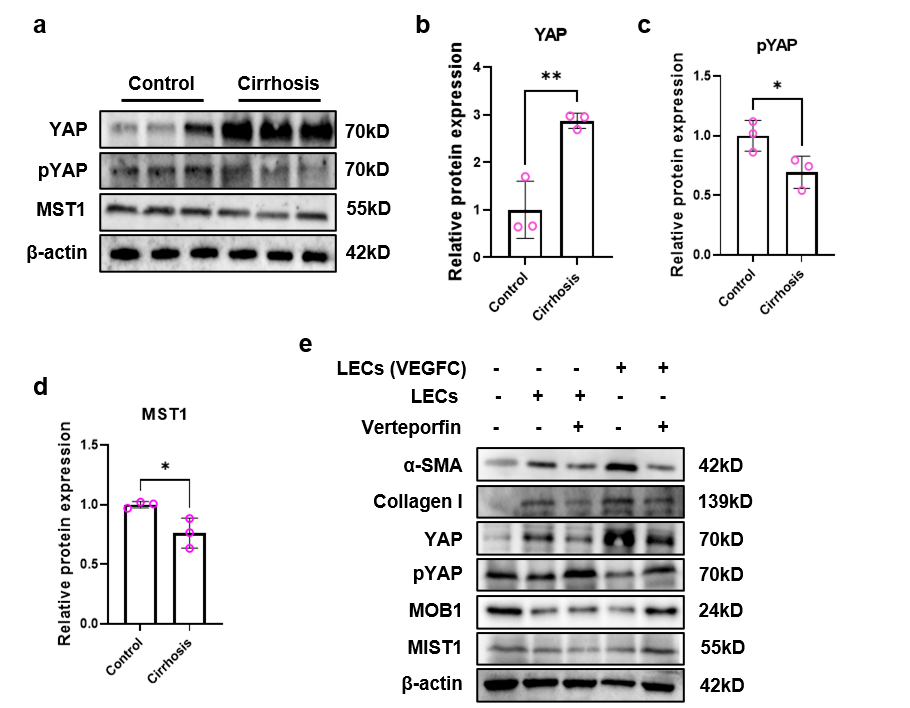

**Fig. S10** Hippo/YAP pathway is involved in LECs-induced HSCs activation. **a-d** Hippo/YAP pathway was detected by immunoblotting in human cirrhotic liver. Quantification was conducted, normalized to β-actin. **e** Activation and Hippo/YAP signaling pathway of LX-2 was measured by immunoblotting.

| **Table S1 Primer sequences** | | |
| --- | --- | --- |
|  | Forward | Reverse |
| Mouse |  |  |
| *rps18* | 5’-TGGGAAGTACAGCCAGGTTC-3’ | 5’-GCAAAGGCCCAGAGACTCAT-3’ |
| *vegfc* | 5’-AGCAGCCCACCCTCAATACC-3’ | 5’-TCCCCACATCTATACACACCTCAC-3’ |
| *il6* | 5’-GAGAGGAGACTTCACAGAGGATACC-3’ | 5’-TCATTTCCACGATTTCCCAGAGAAC-3’ |
| *il1β* | 5’-TCGCAGCAGCACATCAACAAG-3’ | 5’-TCCACGGGAAAGACACAGGTAG-3’ |
| *col1a1* | 5’-GAGCGGAGAGTACTGGATCG-3’ | 5’-GCTTCTTTTCCTTGGGGTTC-3’ |
| *mmp2* | 5’-ACCATGCGGAAGCCAAGATGTG-3’ | 5’-AGGGTCCAGGTCAGGTGTGTAAC-3’ |
| Human |  |  |
| *cd80* | 5’-TCGCCTCTCTGAAGATTACCC-3’ | 5’-CCCAAGTAAGACCAGGGCAC-3’ |
| *cd86* | 5’-GCAGAAGCAGCCAAAATGGAT-3’ | 5’-GACTGAAGTTAGCAGAGAGCA-3’ |
| *cd206* | 5’-ATCACGAAGCCAAGGTCCAG-3’ | 5’-GTGGGTGAACCGAACCTCTT-3’ |
| *vegfc* | 5’-GCTTCTTCTCTGTGGCGTGT-3’ | 5’-CCCGCCTCTCCAAAAAGCTA-3’ |
| *il10* | 5’-AGCTGCTGCCTTGATTGTATT-3’ | 5’-CGTGTGGGTTCAGCCTAGAT-3’ |
| *rps18* | 5’-ATGCAGAATCCACGCCAGTACAAG-3’ | 5’-TCAGTCGCTCCAGGTCTTCACG-3’ |
| *il6* | 5’-CCAGAGCTGTCCAGATGAGTA-3’ | 5’-TGACCTGCCCATGCTACA-3’ |

| **Table S2. Comparison of clinical characteristics in liver fibrosis patients with high versus low serum VEGFC** | | | |
| --- | --- | --- | --- |
| **Clinical features** | **Serum VEGFC** | | **P value** |
|  | Low | High |  |
| Age, mean±SD | 49.25 ± 12.07 | 46.172 ± 14.753 | 0.393 |
| Sex, number (%) |  |  | 0.127 |
| Male | 20 (35.1%) | 15 (26.3%) |  |
| Female | 8 (14%) | 14 (24.6%) |  |
| AST, median | 36 (29.75, 63.5) | 98 (55, 125) | 0 |
| ALT, median | 34.15 (20.7, 55.5) | 89 (48, 109) | 0.005 |
| γ-GGT, median | 91.95 (62.5, 142.68) | 99 (62, 120) | 0.643 |
| ALP, median | 38.5 (29.25, 60) | 39 (24, 66) | 0.962 |
| BAs, median | 6.8 (4, 8.825) | 12.6 (7.1, 19.4) | 0 |
| ADA, median | 1.325 (0.7525, 3.925) | 1.9 (0.6, 5.8) | 0.755 |
| TBIL, median | 23.8 (14.893, 30.525) | 26.6 (12.55, 42.6) | 0.473 |
| TP, median | 70 (65.8, 74.25) | 66 (55, 72) | 0.102 |
| ALB, median | 42 (33.775, 45.25) | 37 (26, 45) | 0.337 |
| TT, median | 27.675 ± 1.9434 | 27.324 ± 1.7537 | 0.477 |
| APTT, median | 9.15 (4.615, 13) | 8.7 (3.65, 11.6) | 0.288 |
| PT, mean±SD | 11.199 ± 4.1499 | 11.783 ± 3.8749 | 0.585 |
| D-D, mean±SD | 11.089 ± 1.4731 | 10.659 ± 1.3824 | 0.26 |
| FIB, median | 0.87 (0.64, 0.9425) | 0.67 (0.36, 0.87) | 0.127 |
| HDL, mean±SD | 1.21 (1.0175, 1.2725) | 1.07 (0.9, 1.22) | 0.079 |
| LDL, mean±SD | 2.4657 ± 0.35127 | 2.8955 ± 0.44272 | 0 |
| **Fibrosis stage, number (%)** |  |  | 0.049 |
| S1 | 5 (8.8%) | 0 (0%) |  |
| S2 | 4 (7%) | 5 (8.8%) |  |
| S3 | 13 (22.8%) | 11 (19.3%) |  |
| S4 | 6 (10.5%) | 13 (22.8%) |  |
| ^AST, aspartate aminotransferase; ALT, alanine aminotransferas; γ-GGT, γ-gamma-glutamyl transferase; ALP, alkaline phosphatase; BAs, bile acids; ADA, adenosine deaminase; TBIL, total bilirubin; TP, total protein; ALB, albumin; TT, thrombin time; APTT, activated partial thromboplastin time; PT, prothrombin time; D-D, D-dimer; FIB, fibrinogen; HDL, high density lipoprotein; LDL, low density lipoprotein.^ | | | |

| **Table S3. Comparison of clinical characteristics in liver fibrosis patients with high versus low serum MDK** | | | |
| --- | --- | --- | --- |
| **Clinical features** | **Serum MDK** | | **P** |
|  | **Low** | **High** |  |
| Age, mean±SD | 41.607 ± 14.415 | 53.552 ± 9.4853 | 0.001 |
| Sex, number (%) |  |  | 0.082 |
| Male | 14 (24.6%) | 21 (36.8%) |  |
| Female | 14 (24.6%) | 8 (14%) |  |
| AST, median | 50.85 (31.15, 86.025) | 87 (37, 125) | 0.133 |
| ALT, median | 41.95 (22.95, 69.75) | 88 (32, 131) | 0.045 |
| γ-GGT, median | 69 (54, 90) | 118 (96, 154) | 0 |
| ALP, median | 34 (26.25, 39.75) | 52 (30, 66) | 0.042 |
| BAs, median | 8.65 (6.5, 11.1) | 8.9 (6.1, 19.4) | 0.363 |
| ADA, median | 0.845 (0.3, 1.825) | 4.7 (1.33, 7.7) | 0.001 |
| TBIL, median | 16.75 (10.4, 26.95) | 27.1 (22, 40.8) | 0.001 |
| TP, median | 70.1 (67, 74) | 65 (53, 71) | 0.021 |
| ALB, median | 42.55 (37.375, 46.225) | 31 (24, 44) | 0.001 |
| TT, mean±SD | 27.675 ± 2.0038 | 27.324 ± 1.6869 | 0.477 |
| APTT, median | 10.75 (8.375, 13.825) | 4.65 (3.41, 9.1) | 0.001 |
| PT, mean±SD | 11.006 ± 4.5047 | 11.969 ± 3.4287 | 0.367 |
| D-D, mean±SD | 10.825 ± 1.5997 | 10.914 ± 1.2752 | 0.817 |
| FIB, mean±SD | 0.67 (0.33, 0.87) | 0.85 (0.65, 0.97) | 0.098 |
| HDL, median | 1.21 (1.06, 1.275) | 1.01 (0.9, 1.24) | 0.037 |
| LDL, median | 2.5 (2.3525, 2.825) | 2.84 (2.4, 2.99) | 0.157 |
| **Fibrosis stage, number(%)** |  |  | 0 |
| S1 | 5 (8.8%) | 0 (0%) |  |
| S2 | 8 (14%) | 1 (1.8%) |  |
| S3 | 14 (24.6%) | 10 (17.5%) |  |
| S4 | 1 (1.8%) | 18 (31.6%) |  |
